# Amplification-Free Dual-Blocking Autocatalytic CRISPR-Cascade for Atto-Molar DNA Detection with Low Nonspecific Signal

**DOI:** 10.64898/2025.12.03.692097

**Authors:** Jongwon Lim, An Bao Van, Matthew Wester, Katherine Koprowski, Enrique Valera, Rashid Bashir

## Abstract

Autocatalytic CRISPR architecture offers amplification-free nucleic acid detection by directly linking target recognition to self-reinforcing ribonucleoprotein (RNP) generation. However, spontaneous background activation remains a key barrier, because strand invasion or unwinding events can initiate unintended amplification and diminish assay specificity. Here, we introduce a dual-blocking CRISPR-Cascade design that independently cages both the guide RNA and trigger DNA, establishing an intrinsic AND gate to raise the effective kinetic barrier for unintended RNP formation. This strategy suppresses leakage by approximately 3- to 18-fold relative to single blocking configurations in full Cascade reactions, while preserving rapid detection (10 min), achieving single-copy sensitivity, and enabling quantitative detection. When paired with a competitive guide RNA decoy, the system further reduces background signals without affecting true target detection. Finally, we demonstrate robust Methicillin-resistant Staphylococcus aureus (MRSA) detection from whole blood in under 40 minutes including the sample purification and extraction. These results establish dual-blocking as a generalizable molecular gating framework for constructing leakage-resistant, amplification-free CRISPR systems suitable for rapid and decentralized diagnostics.

**Significance:** Amplification-free CRISPR diagnostics are often presented as simple positive feedback circuits, but most existing systems treat leakage, defined as target independent background activation that arises when blocked CRISPR components spontaneously form active ribonucleoprotein (RNP), as an unavoidable side effect rather than as a designable property. In particular, prior work has not explicitly accounted for two key sources of background signal in autocatalytic assays: Cas driven unwinding of blocked constructs and transient breathing of nucleic acid duplexes that intermittently expose trigger sites. Our study directly analyzes these leakage pathways in the context of switchable-cage-gRNA (scgRNA) and Cascade probe design and shows that blocking a single component is fundamentally vulnerable to both enzyme-driven strand invasion and equilibrium breathing. By contrast, we introduce a dual-blocking strategy in which both the guide RNA and the trigger DNA are gated. We further add a decoy guide RNA that competes for Cas12a. This multi-layer architecture demonstrates that robust amplification-free operation requires several coordinated barriers rather than a single switch, providing a new design principle for constructing self-amplifying CRISPR circuits with low background and robust signal-to-noise ratio.

## Introduction

Nucleic acid diagnostics traditionally rely on polymerase chain reaction (PCR) or other enzymatic amplification methods to achieve high sensitivity.^1,2^ Although highly effective, these workflows require multistep sample processing, precise temperature control, and pose a risk of carryover contamination, making them less suitable for rapid or decentralized testing.^3^ CRISPR-based detection systems have addressed several of these limitations by offering programmable target recognition and operation under isothermal reaction conditions.^4,5^ However, most CRISPR-based assays still require an upstream nucleic acid amplification step to achieve clinically relevant levels of detection sensitivity.^6–8^

To eliminate this dependency, autocatalytic CRISPR architectures have recently emerged as a strategy to couple target recognition directly to self-reinforcing ribonucleoprotein (RNP) activation.^9^ In these systems, the key molecules are blocked nucleic acids (BNAs), which are switchable intermediates or blocked precursors. An initial target-activated Cas complex converts inactive BNAs into new active molecules, forming a positive feedback loop that enables exponential signal gain without thermal cycling.^10–12^ Several BNA designs have been developed to achieve conditional activation, including cleavage-activated guide-releasing constructs,^9^ intramolecularly folded trigger strands stabilized by nucleic-acid chemistries,^13,14^ duplex-protected DNA/RNA triggers,^15^ and topology-gated circular nucleic acids that require linearization.^16^ Collectively, these approaches demonstrate that nucleic-acid structures can be programmed to serve as feedback gates that drive self-catalyzing CRISPR reaction.

Despite these advances, all existing autocatalytic CRISPR systems share a fundamental challenge: spontaneous background activation.^11,12^ Transient breathing or equilibrium fluctuations of BNA, or strand-invasion-driven unwinding by Cas enzymes, can generate small amounts of active RNPs that propagate through the positive feedback loop. This leakage causes fluorescence accumulation in the absence of target, narrows the dynamic range, limits quantification, complicates thresholding, and ultimately limits confidence at low target concentrations, precisely where amplification-free systems offer the greatest potential impact.

To overcome this limitation, we developed a dual-blocking architecture that simultaneously cages the guide RNA (gRNA) and the trigger DNA required to form the amplifying Cas12a complex. Productive RNP assembly, and thus initiation of the feedback loop, requires coordinated release of both components, enforcing an intrinsic AND-gate logic at the core of autocatalytic activation. This design establishes two orthogonal kinetic barriers against unintended RNP formation, substantially elevating the activation threshold relative to single-blocking systems. To further suppress rare leakage events, we also introduce a sacrificial guide RNA decoy that competitively captures free Cas enzymes before they can engage the blocked substrates. Together, these strategies lower the negative baseline signal while preserving rapid exponential amplification upon target recognition, enabling sensitive, amplification-free detection with greatly reduced false activation risk. Using this architecture, we demonstrate rapid and specific detection of MRSA, including successful identification of low-copy targets obtained through extraction and purification from whole blood samples.

## Results

### Rationale for dual-blocking of autocatalytic CRISPR-Cascade reaction systems

Autocatalytic CRISPR reactions enable highly sensitive detection but are often compromised by spontaneous background amplification caused by unintended activation of the positive feedback loop. The CRISPR-Cascade reaction is initiated when RNP T1, Cas12a pre-complexed with a target-specific gRNA, recognizes and cleaves target DNA in cis, and is activated as a relatively indiscriminate trans-nuclease (Fig. 1a). Activated RNP T1 then triggers the downstream autocatalytic reaction by converting latent precursors into active components of RNP T2.

**Figure 1.**
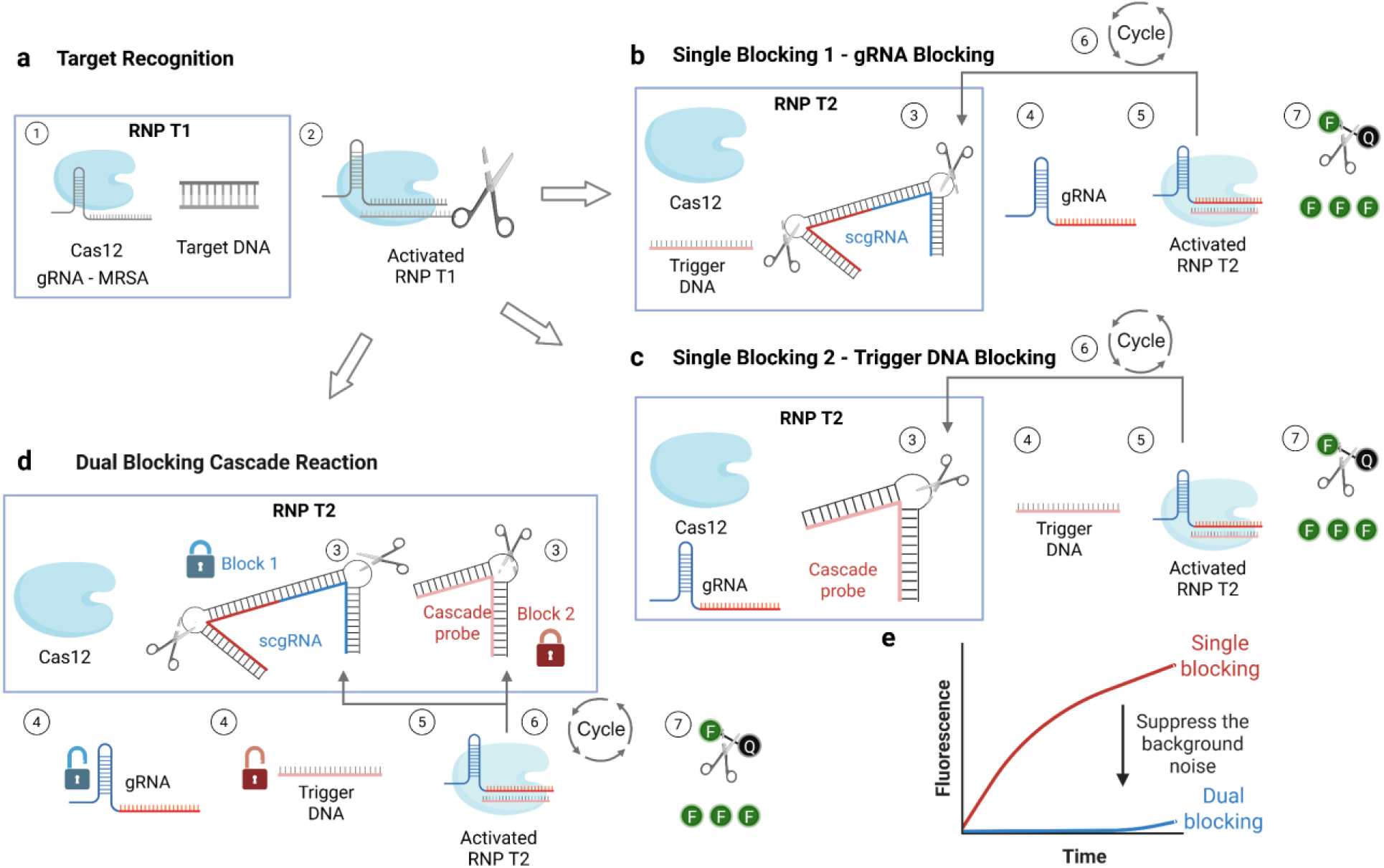
Dual-blocking strategy for suppressing background signals in autocatalytic CRISPR cascade reactions. The Cascade mechanism is modularized into (a) primary target recognition (RNP T1) and (b–d) downstream signal amplification (RNP T2). (a) RNP T1 becomes activated upon binding and cleaving target DNA. (b–c) In single-blocking configurations, either the switchable caged guide RNA (scgRNA for gRNA-blocking) or Cascade probe (trigger-blocking) protects one precursor. Activated RNP T1 cleaves the bulge structures to release either gRNA or trigger DNA, enabling RNP T2 formation and self-amplifying positive feedback. (d) In dual-blocking, both precursors are caged, requiring two independent unlock steps before RNP T2 assembly, thus elevating the activation threshold and limiting leakage. (e) Dual-blocking markedly reduces spontaneous background fluorescence compared to single blocking.

Previous BNA designs have relied on a single-blocking strategy in which only one precursor species was protected to create a switchable intermediate. Specifically, a switchable caged guide RNA (scgRNA, Fig. 1b) and Cascade probe (Fig. 1c) are pre-assembled by blocking the gRNA and trigger DNA, respectively, and are supplied in the RNP T2 tube as switchable precursors. In both approaches, engineered single-stranded DNA (ssDNA) bulges act as conditional gates; cleavage of the bulge by activated RNP T1 releases the guide RNA or trigger DNA, which in turn increases the number of activated RNP T2 and reinforces the autocatalytic loop. Here, it is important to note that the corresponding unprotected strand (e.g., trigger DNA for scgRNA systems and gRNA for Cascade-probe systems) is provided in high concentration (tens to hundreds of nanomolar) in the RNP T2 mixture. Consequently, once the trans-cleavage activity of activated RNP T1 cleaves the BNA bulges and releases small amounts of active gRNA or trigger DNA, RNP T2 complexes form rapidly, thereby initiating and amplifying the autocatalytic feedback loop.

However, single-blocking systems often fail to suppress nonspecific background signals due to two intrinsic leakage mechanisms arising from: (i) Cas enzyme–driven strand invasion and (ii) transient base-pair “breathing” of blocked BNA duplexes at thermodynamic equilibrium. In the first case, Cas12a can spontaneously invade and open scgRNA to form functional RNP complexes, or preassembled RNPs can intermittently strand-invade and unwind Cascade probes, transiently activating the feedback loop through unintended RNP T2 formation (Fig. S1a–S1b).^17,18^ In the second case, the coexistence of unlocked and blocked BNA species at thermodynamic equilibrium allows partial exposure of the recognition site, promoting stochastic initiation events (Fig. S1c–S1d).^19,20^ Together, these two mechanisms synergistically contribute to background nonspecific signals in single-blocking systems. A detailed theoretical justification for this behavior, including the derivation of how thermodynamic leakage scales with the equilibrium bound fraction α and why dual-blocking yields multiplicative suppression of the respiratory leakage term, can be found in Supplementary Note 2.

To minimize these leakage effects, we developed a dual-blocking system that simultaneously cages both the gRNA and trigger DNA using the scgRNA and Cascade probe (Fig. 1d). Upon target recognition, RNP T1 cleaves the ssDNA bulges in both constructs, releasing unblocked gRNA and trigger DNA, and enabling productive formation of RNP T2.^13^ This architecture therefore requires both BNA components to be activated, implementing an AND-logic gate, and imposing two orthogonal kinetic barriers to spontaneous activation by independently caging both gRNA and trigger DNA (Fig. S1e). As a result, the dual-blocking architecture markedly suppresses background noise compared to the single-blocking approach (Fig. 1e).

### Construction and baseline evaluation of dual-blocking components

To construct the dual-blocking system, we first adapted a scgRNA architecture in which the gRNA is hybridized with two inhibitor DNAs, iDNA-handle (iH) and iDNA-spacer (iS), following the design of Shi *et al* (Fig. 2a).^9^ The spacer region of gRNA was used to define the trigger DNA (TD) sequence, and its complementary strand was engineered into a bubble DNA (BD) containing a seven-base single-stranded bulge (TTTATTT) at the middle location of bubble DNA to generate a Cascade probe that cages the trigger DNA (Fig. 2b). Thermodynamic predictions from NUPACK^21^ and DINAMelt^22^ indicated stable secondary structures for both scgRNA and the Cascade probe (Fig. S2a - S2c). Experimental validation using melting-curve analysis and native 15% PAGE confirmed correct formation of both blocked complexes (scgRNA and CP) from individual components (iH, iS, gRNA, TD, BD), consistent with simulation output (Fig. 2c and 2d).

**Figure 2.**
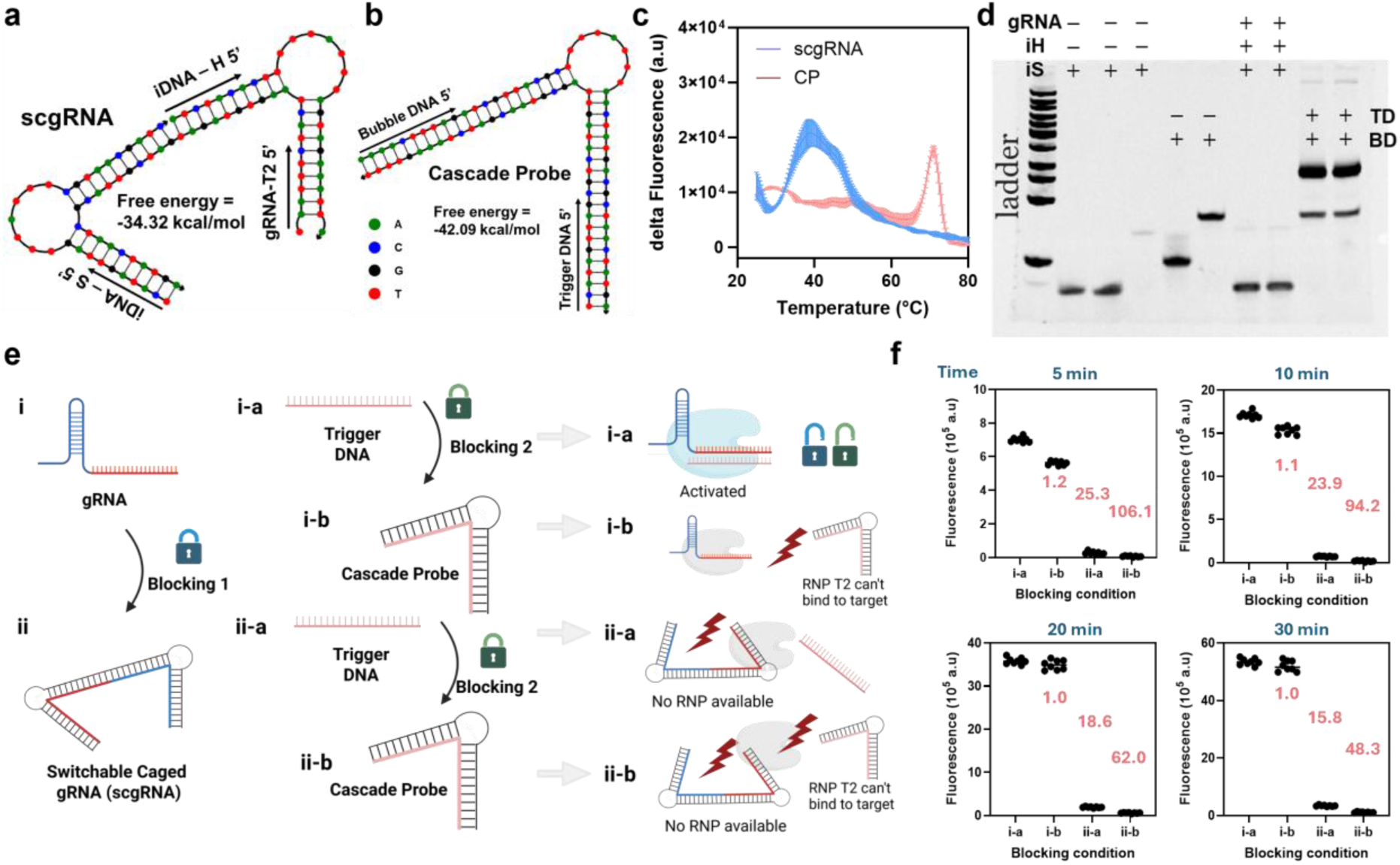
Construction and baseline evaluation of dual-blocking components. (a–b) NUPACK predicted secondary structures of scgRNA and the Cascade probe, with annotated 5′ starting points for gRNA-T2, inhibitor handle (iH), inhibitor spacer (iS), trigger DNA (TD), and bubble DNA (BD), along with their calculated free energies. (c) Melting-curve profiles of assembled scgRNA and Cascade probe. (d) Native PAGE analysis of scgRNA and Cascade probe alongside individual component strands. (e) A 2×2 single Cas12a reaction matrix was used to isolate RNP T2 formation and evaluate four precursor states: unblocked (i-a), trigger-blocked (i-b), gRNA-blocked (ii-a), and dual-blocked (ii-b). (f) Fluorescence signal at various time points of the four precursor states in the single-Cas12a assay. Signal reduction (fold), defined as the fluorescence ratio relative to the unblocked state, are shown in red for each time-point and condition. Data are presented as mean ± error (n = 3) in panel c, and as individual replicate traces (n = 8) in panel f.

To quantify the blocking effect of BNA independent from upstream activation, we implemented a single-Cas12a assay that isolates RNP T2 formation. In this configuration, either the gRNA or trigger DNA (or both) remained blocked, generating a 2×2 matrix with four states: unblocked (i-a), trigger DNA-blocked (i-b), gRNA-blocked (ii-a), and dual-blocked (ii-b) (Fig. 2e). Fluorescence kinetics revealed progressive signal reduction as each blocking layer was introduced, with the lowest signal observed in the dual-blocked state (Fig. 2f, Fig. S2d and S2e). Blocking efficiencies (or signal reduction), calculated as the fluorescence ratio of the fully unblocked condition (i-a) to each blocked state, demonstrated that scgRNA alone provided ∼23.9-fold suppression at 10 min, whereas the Cascade probe alone yielded only ∼1.1-fold suppression. Strikingly, dual-blocking achieved ∼94.2-fold suppression, with the strongest effect at early timepoints (∼106.1-fold at 5 min), suggesting efficient prevention of premature RNP formation that would otherwise trigger the positive-feedback reaction (Fig. S1). This suppression exceeded the expected product of the individual blocking efficiencies, indicating a synergistic interaction between the orthogonal gRNA- and trigger-blocking pathways.

These single-Cas12a results using RNP T2 components show not only the advantage of combining scgRNA and the Cascade probe but also help us to understand why existing CRISPR autocatalytic systems often display substantial background leakage. When transitioning from the unblocked to trigger-blocked condition (i-a to i-b), signal persisted (Fig. 2f and S2d) because the RNP can easily strand-invade the Cascade probe and partially unwind the duplex, thereby exposing the target regions of trigger DNA and enabling activation even in the blocked state (Fig. S1b).^17,18^ Conversely, transitioning from unblocked to gRNA-blocked (i-a to ii-a) revealed that Cas12a can also invade scgRNA and form functional RNP despite inhibitory strands (Fig. S1a). Surprisingly, even when both components were blocked (ii-b), signal gradually emerged over time (blue line, Fig. S2e). If this background originated solely from unpaired or incompletely blocked DNA (thermodynamic respiration), providing excess complementary blockers should have strongly suppressed the signal. However, even increasing the blocker ratios (gRNA:iDNAs and trigger:bubble) from 1:1 up to 1:2 only modestly reduced leakage and did not fully eliminate it (Fig. S2f and S2g). This observation suggests that the background activity cannot be explained by equilibrium dynamics alone but instead reflects a kinetic bypass pathway in which Cas12a could actively invade and unwind apparently stable blocking structures, leading to residual activation despite strengthened thermodynamic pairing. A quantitative description of this leakage suppression trend as a function of blocker stoichiometry (x, y, z) and equilibrium parameters is provided in the Supplementary Note 1 (Section “varying complement concentration”).

Finally, we verified that target-activated RNP T1 can cleave ssDNA bulges and release functional products (Fig. S3). Incubation of the scgRNA or Cascade probe with the activated RNP T1 (detailed protocol in Fig. S3a) generated trans-cleavage fragments (Fig. S3b and S3d). Increasing pre-cleavage time led to progressively higher downstream signals in reporter assays (Fig. S3c and S3e), confirming that unblocked gRNA and trigger DNA produced by activated RNP T1 serve as functional RNPs for downstream activation. Together, these data validate the dual-blocking module architecture, quantify its suppression capacity, and provide a potential mechanistic explanation for leakage in prior CRISPR autocatalytic systems which underscores the necessity of an orthogonal two-barrier strategy (Fig. S1d).

### Engineering Cascade probe stability to balance dual-blocking contributions

While dual-blocking substantially reduced background compared to the single-blocking strategy (Fig. 2f), the two blocking modules did not contribute equally to leakage suppression. For example, the gRNA-blocking module (scgRNA) imposed a much stronger barrier (25.3-fold suppression at 5 min, Fig. 2f), whereas the trigger-blocking module (Cascade probe) offered only modest suppression in its initial configuration (1.2-fold suppression at 5 min, Fig. 2f), resulting in an imbalanced AND-gate behavior (Fig. 3a). Although the imbalance itself is not inherently detrimental, it indicates that the trigger-blocking pathway offers greater room for optimization compared to the gRNA-blocking module. To strengthen the trigger-blocking pathway, we systematically engineered Cascade probe designs by varying: (i) bubble-DNA length for partially blocked Cascade probes, (ii) the number of bulge units embedded in the bubble DNA, (iii) placement of locked nucleic acids (LNAs) adjacent to bulges to enhance local stability,^23,24^ (iv) segmentation of the bubble DNA into two parts, analogous to scgRNA’s iDNAs,^9^ and (v) the trigger-DNA sequence (Fig. 3b and Figs. S4–S5).

**Figure 3.**
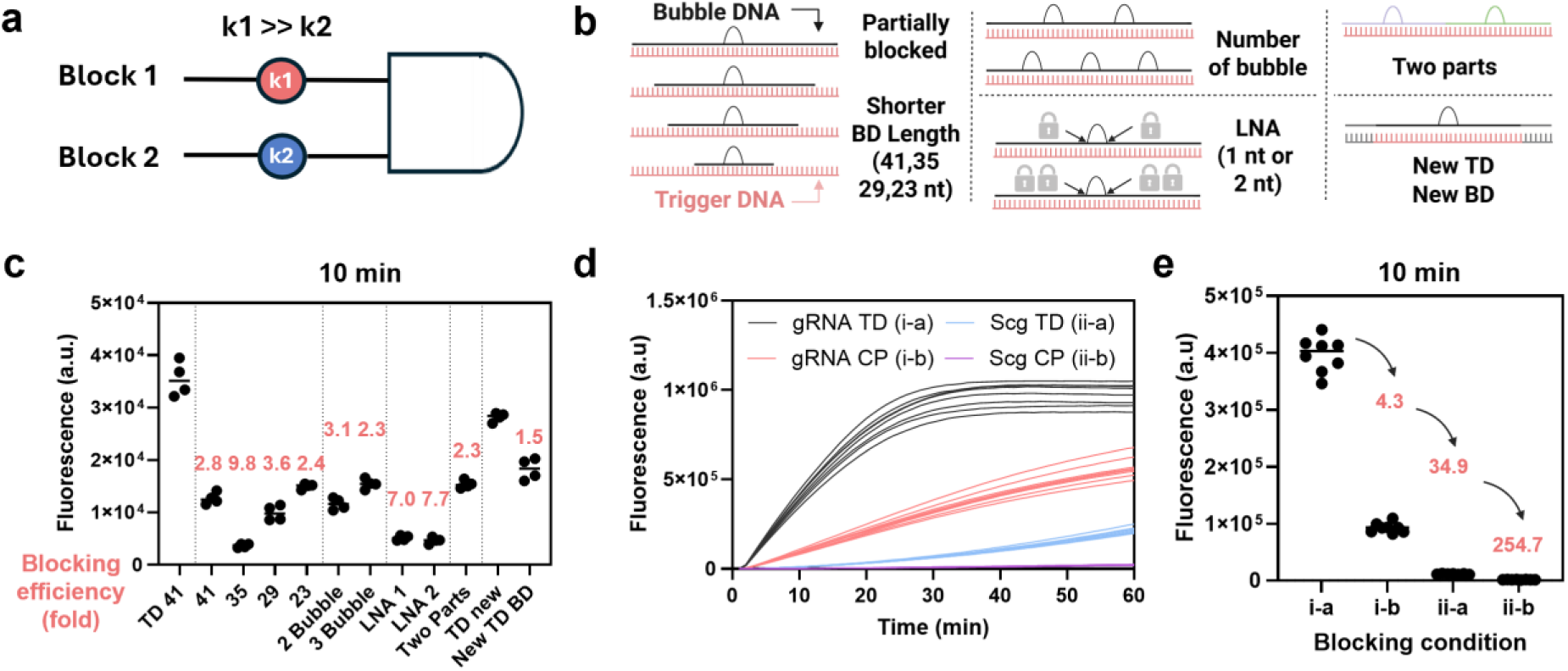
Engineering Cascade probe stability to balance dual-blocking contributions. (a) Schematic illustrating the initial asymmetry between gRNA-blocking (scgRNA) and trigger-blocking (Cascade probe) pathways in the dual-blocking system. The scgRNA contributed a disproportionately stronger inhibition relative to the Cascade probe (k₁ >> k₂), resulting in an unbalanced AND-gate behavior. (b) Cascade probe variants tested, including designs with altered bubble-DNA length, bulge count, LNA placement, duplex segmentation, and trigger-DNA sequence. (c) Single-Cas12a assay comparing fluorescence profiles at t=10 minutes using gRNA with either unblocked trigger DNA (“TD41” or “TD-new”) or Cascade probe variants. Bubble-DNA lengths (41, 35, 29, and 23 nt) represent designs shortened in 6-nt increments (excluding bulges); “2-bubble” and “3-bubble” indicate the number of bulge units; “LNA-1” and “LNA-2” indicate incorporation of one or two LNAs beneath the bulge; “two-parts” indicates division of the bubble-DNA into two segments; “TD-new” denotes an alternative trigger-DNA sequence; and “new-TD-BD” is the corresponding Cascade probe for that trigger. Blocking efficiencies are shown in red. For example, the TD41 signal divided by the CP41 signal yields a blocking efficiency of 2.8. (d) Representative 2×2 single-Cas12a kinetic traces using the LNA-1 probe variant. (e) End-point signal and calculated blocking efficiencies using the LNA-1 probe. Data are shown with individual replicates (n = 4 for panel c, and n = 8 for panels d and e).

Single-Cas12a measurements using gRNA together with either trigger DNA (TD41) or various Cascade probes (corresponding to the i-a and i-b conditions in Fig. 2f) showed that Cascade probes containing a 35-nt bubble DNA (BD35) or LNA-modified bulges achieved the strongest blocking, with 7.0–9.8-fold suppression relative to unblocked TD41 controls (Fig. 3c). Representative kinetic profiles using the LNA-optimized Cascade probe confirmed that the contribution of the two blocking modules became more balanced (Fig. 3d). In the unblocked condition (i-a), signal rose rapidly as expected. When only the Cascade module was blocked (i-b), suppression reached 4.3-fold, whereas scgRNA-only blocking (ii-a) yielded 34.9-fold suppression at 10 minutes (Fig. 3e). Dual-blocking (ii-b) resulted in the strongest suppression, giving ∼254.7-fold signal reduction. This combined effect exceeded the expected product of the individual-blocking efficiencies by approximately 1.7-fold, indicating a measurable synergistic interaction between the two blocking modules. Full datasets at 10, 20, and 30 minutes are provided in Figs. S6–S7. These results show that increasing the blocking efficiency of the Cascade probe narrows the performance gap between the two blocking pathways and produces a more balanced AND-gate response. However, blocking efficiency alone does not always correlate with robust switching performance, emphasizing the need for full assay characterization.

Regardless of blocking-probe enhancements, scgRNA blocking remained the dominant contributor to suppression, even under optimized probe designs. This behavior reflects the intrinsic reaction order in which Cas12a must first bind gRNA to form the initial RNP complex before engaging any trigger substrate.^25,26^ Accordingly, gRNA sequestration among the two BNA components imposes the primary kinetic checkpoint, yielding a sequential AND-gate architecture initiated by scgRNA (Fig. S8).

### Enabling ultrasensitive CRISPR detection by integrating dual-blocking into the full Cascade workflow

Next, we implemented the dual-blocking architecture within a complete CRISPR-Cascade assay. To enable systematic characterization of the assay, we first defined a consistent reference configuration. Two sample inputs were used throughout: MRSA genomic DNA (10⁴ copies/µL) as the positive control and a no-template negative control, both in nuclease-free water. For SNR calculations, these positive and negative samples were measured in parallel under identical conditions. The concentrations of the two blocking modules, scgRNA and Cascade probe, were fixed at 10 nM each (Fig. S9), and the RNP T1:RNP T2 ratio was set to 1:3, using CP-41 as the reference Cascade probe. This baseline served as the reference condition for all subsequent optimization and benchmarking studies, unless otherwise specified for configurations yielding superior performance. Under this condition, we compared the signal-to-noise ratio (SNR) between positive and negative samples at t = 10 min, establishing a quantitative reference point for subsequent performance improvements.

We first optimized the reaction sequence. Cas12a was first pre-complexed with MRSA-targeting gRNA (pre-incubation, 5 min; Fig. 4a-i), followed by target DNA activation (pre-activation, 5 min; Fig. 4a-ii). The activated RNP T1 complex was then introduced to the dual-blocked T2 components (Fig. 4a-iii), enabling RNP T1 to turn on the autocatalytic switch and RNP T2 to drive positive-feedback amplification (Fig. 4b). To identify the most effective reaction workflow, we evaluated three configurations (Fig. 4c): (i) pre activation only, in which Cas12a, gRNA, and target DNA were introduced simultaneously without prior RNP assembly (“i ii combined”); (ii) sequential pre incubation followed by pre activation, allowing RNP T1 formation before target engagement (“Final Protocol”); and (iii) one pot assembly, in which all components, including the T2 module, were combined at the outset with no pre complex formation (“i ii iii combined”). The sequential protocol (ii) integrating pre-incubation and pre-activation produced the strongest signal separation between positive and negative samples, demonstrating that staged RNP assembly and activation enhances controlled engagement of the T2 autocatalytic module.

**Figure 4.**
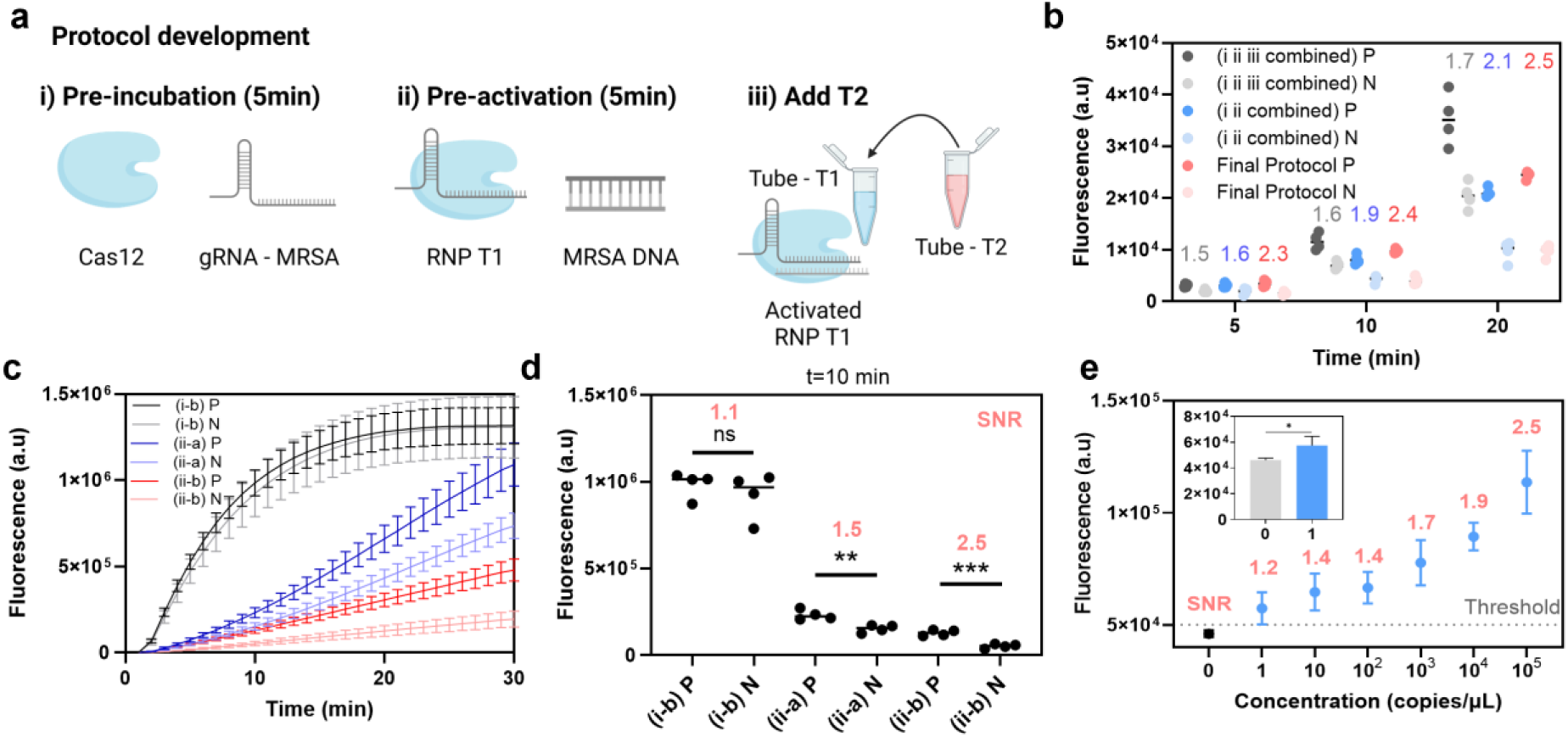
Integrating dual-blocking into the full CRISPR Cascade workflow. (a) Schematic of the reaction sequence: (i) Cas12a pre-incubation with gRNA-MRSA forming RNP complex, (ii) target DNA pre-activation by adding sample, and (iii) introduction of the dual-blocked RNP T2 components into the activated RNP T1. (b) Comparison of three reaction workflows: pre activation without pre incubation ((i ii combined), blue), sequential pre incubation followed by pre activation (Final Protocol, red), and one pot assembly of all components ((i ii iii combined), gray). End point fluorescence is shown at 5, 10, and 20 minutes. Positive samples, denoted as P (10⁴ copies per microliter MRSA genomic DNA), are shown in darker colors, and negative samples, denoted as N (no template), are shown in lighter colors. The SNR for each case is displayed numerically in the corresponding color. (c) Kinetic traces for positive samples (P, shown in darker colors) and negative samples (N, shown in lighter colors) under dual-blocking and single blocking configurations. The condition using scgRNA with trigger DNA (scg TD, blue) represents single blocking 1, the condition using gRNA with Cascade probe (gRNA CP, red) represents single blocking 2, and the condition using both scgRNA and Cascade probe (scg CP, gray) represents dual-blocking. (d) End-point signal and SNR analysis at 10 minutes demonstrates that dual-blocking improves discrimination by disproportionately suppressing leakage. (e) Serial dilution of MRSA genomic DNA showing end-point readout at 10 minutes from single copy to 10⁴ copies per microliter. The signal-to-noise ratio (SNR) was calculated by dividing each concentration by the corresponding negative sample and is displayed in red. The inset shows statistical analysis comparing negative and lowest positive (1 copy per microliter) samples from the full calibration. Statistical analysis was performed using a two tailed unpaired t-test; asterisk indicates P less than 0.05. Data are presented with individual replicates (n = 4 for panel b and d), and with mean and standard deviation (n = 4 for panel c and n = 5 for panel e).

Using this optimized workflow, we then evaluated assay performance under varied molecular designs and reaction conditions. First, we screened different Cascade probe variants. Surprisingly, CP29, rather than CP35 or the LNA-modified probe, yielded the highest SNR (1.87; Fig. S10), demonstrating that maximal blocking efficiency (Fig. 3c) does not necessarily translate to optimal switching fidelity or assay accuracy. This is because blocking strength reduces background but does not dictate how quickly RNP T1 can cleave the bulge and amplify signal, making SNR dependent on the balance between kinetic activation and background suppression rather than blocking efficiency alone. Guided by this result, we used CP-29 as the reference probe for subsequent optimization. We next evaluated buffer composition (Fig. S11a) and RNP T1:T2 ratios (Fig. S11b and S11c). Together, these analyses identified the optimal configuration as 10 nM scgRNA, 10 nM Cascade probe, NEB Buffer 2.1, a 1:3 RNP T1:T2 ratio (corresponding to a final concentration of 6 nM for both scgRNA and Cascade probe), and CP-29 as the standard Cascade probe for subsequent experiments.

We next benchmarked the dual-blocking circuit against each single-blocking configuration (Fig. 4b and Fig. 4d). All reaction conditions were identical across groups, except that one blocking element (scgRNA or Cascade probe) was replaced with its unblocked counterpart (gRNA or trigger DNA) at the same concentration to ensure a fair comparison. Dual-blocking markedly suppressed background, reducing leakage by 2.94-fold relative to scgRNA-only blocking, and by 17.81-fold relative to trigger-DNA blocking (Fig. S12a). This is consistent with leakage trends observed in the single-Cas12a characterization of RNP T2 (Fig. 2f and Fig. 3e). While positive signals also decreased (7.66-fold and 1.80-fold respectively, Fig. S12b), the overall signal-to-noise ratio (SNR) improved (Fig. S12c and S12d). As a result, scgRNA-only blocking yielded a modest SNR of 1.5 at 10 minutes, and trigger-DNA-only blocking produced a near-baseline SNR of 1.1, whereas the dual-blocking architecture achieved a substantially higher SNR (2.5 at 10 min; Fig. S12c). These data indicate that dual-blocking attenuates spurious activation disproportionately more than it diminishes the true positive signal. For comparison, a single-Cas12a reaction lacking the Cascade signal amplification detected only ∼10 pM targets (Fig. S13), illustrating the intrinsic sensitivity limit without positive feedback, and underscoring that autocatalytic amplification is essential for highly sensitive detection.^27,28^

Finally, 10-fold serial dilutions in nuclease-free water of MRSA genomic DNA revealed detection down to the single-copy regime, with SNR values between 1.2 and 2.5 across replicates (Fig. 4e). Full calibration curves, including titration behavior across the dilution series, are provided in Fig. S14. For the one-pot assembly reaction, the calibration curve showed poor discrimination in fluorescence over time between concentrations below 10³ copies/µL (Fig. S15), highlighting the importance of supplying RNP T1 separately from the autocatalytic RNP T2 layer to fully preserve sensitivity. Together, these results establish dual-blocking as an effective strategy that not only suppresses spontaneous leakage, but also enables attomolar-level detection capability in a self-amplifying CRISPR system.

### Eliminating residual leakage by a sacrificial guide RNA decoy

Although dual-blocking substantially reduced leakage and improved signal discrimination (Fig. 4d), residual background activity remained detectable. To further suppress unintended activation, we implemented a simple competitive strategy in which a large molar excess of free gRNA T1 (up to tens of micromolar) was supplied as sacrificial decoys to outcompete premature RNP T2 formation (Fig. 5a). Because gRNA does not trigger trans-cleavage on its own, it can be supplied in large excess as a decoy without interfering with Cas12a activity or overall reaction kinetics. During the T1 incubation step, Cas12a is already saturated with its guide RNA, so the additional decoy gRNAs remain unbound. These free gRNAs remain inactive until the T2 components are added, where they preferentially associate with unbound Cas12a rather than the blocked scgRNA, effectively diverting the enzyme from premature RNP T2 formation. This diversion effectively reduces unintended activation driven by scgRNA unwinding (Fig. S1a and Fig. S16).

**Figure 5.**
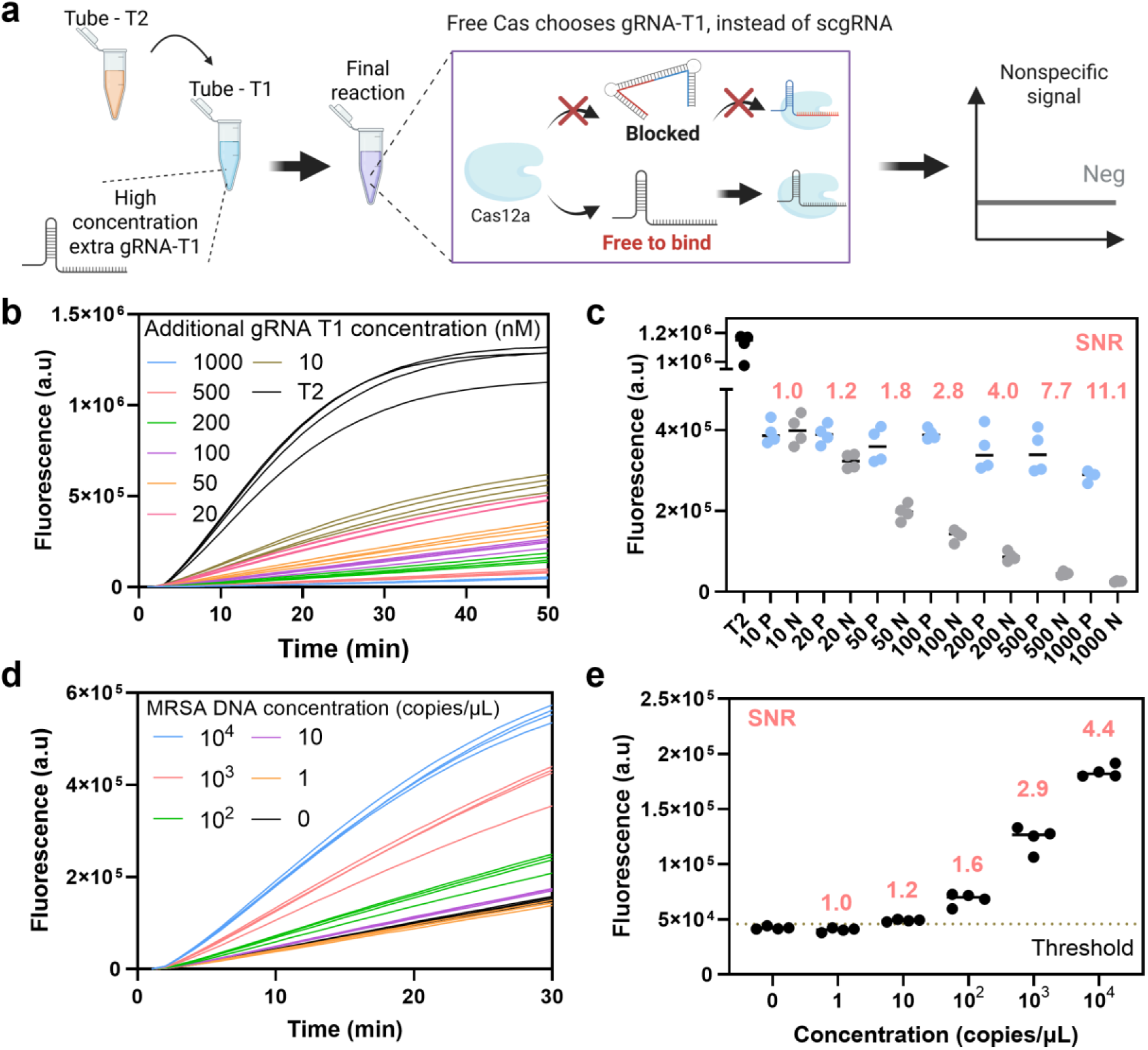
Suppressing residual leakage using a competitive gRNA decoy strategy. (a) Schematic illustrating the decoy concept. Extra gRNA T1 is supplied in T1 tube, remaining unbound until mixed with T2 components. Free Cas12a in the T2 tube preferentially binds the unblocked decoy gRNA T1 rather than the blocked scgRNA. This competitive binding prevents premature RNP T2 formation driven by scgRNA unwinding, resulting in reduced nonspecific signal. (b) Negative control fluorescence traces with increasing concentrations of decoy gRNA T1, showing progressive suppression of background activation. (c) End point fluorescence at 30 minutes for positive (10⁴ copies per microliter MRSA genomic DNA) and negative (no template) are shown at corresponding decoy concentrations (nM), showing minimal effect on true signal generation. The SNR is displayed above each condition with purple color. (d) Detection of MRSA genomic DNA at varying concentrations from 10⁴ to 1 copies per microliter with decoy gRNA protocol. (e) End point signal and signal to noise ratio at 10 minutes across MRSA dilutions. Data is shown with individual well replicates (n = 4 for all panels).

Critically, the decoy strategy selectively suppressed negative-control leakage without compromising true signals. When only the T2 module was present without RNP T1, strong spontaneous activation occurred (black line, Fig. 5b). In the full Cascade, adding increasing amounts of decoy gRNA progressively reduced background amplification under negative-control conditions (Fig. 5b), while positive reactions (10⁴ copies/µL MRSA DNA) remained stable at ∼3–4 × 10⁵ a.u. across all decoy concentrations (Fig. 5c). As a result, the SNR increased from ∼1.0 to ∼11.1 as the decoy concentration was raised from 500 nM to 50 µM. Unlike the addition of background DNA, which reduces both positive and negative signals by diverting Cas12a trans-cleavage activity,^13^ the gRNA-decoy operates through a trans-cleavage-independent mechanism: it selectively sequesters free Cas12a before unintended RNP formation, thereby eliminating leakage without affecting reaction kinetics. In comparison, adding 3 ng/µL human genomic DNA reduced both positive and negative fluorescence (Fig. S17a), yielding only a modest SNR improvement (∼2.58 to ∼3.19; Fig. S17b).

The effect of the decoy gRNA was sequence-independent. Unrelated gRNA sequences used previously by our group produced similar suppression trends (Fig. S18), indicating that the benefit arises from competitive Cas12a binding rather than target-dependent interactions. Extended monitoring showed that the leakage signal remained at baseline for several hours (Fig. S19), reminiscent of PCR reactions, in which true negatives remain flat. Combining multiple MRSA-targeting guides further enhanced the signal output (9.8 to 12.2; Fig. S20), consistent with prior reports of multi-guide synergy in CRISPR systems. Based on these results, we used four MRSA-targeting decoy gRNAs,^29–31^ each at 5 µM, and evaluated quantitative performance across MRSA genomic DNA concentrations from 10⁴ to 1 copies/µL (Fig. 5d and 5e). With the decoy strategy, SNR at 10⁴ copies/µL improved from ∼1.9 (Fig. 4e) to ∼4.4 (Fig. 5d), while ultra-low inputs remained detectable but with slightly reduced sensitivity, slightly raising the practical detection limit from 1 copy per µL to about 10 copies per µL (SNR ≈ 1.2).

### Analysis of blood sample with dual-blocking

Lastly, we evaluated the performance of our dual-blocking CRISPR-Cascade assay using mock clinical samples spiked with MRSA pathogens in whole blood. Whole blood was inoculated with MRSA at 10-fold serial dilutions to simulate clinically relevant pathogen loads, followed by standard lysis and DNA extraction using commercial kits (Fig. 5a). The extracted nucleic acids were then tested using our optimized dual-blocking configuration, which combines CP29, a decoy gRNA-T1, and a four-guide MRSA gRNA-T1 pool.

The total time-to-result was 40 minutes, consisting of 10 minutes for thermal lysis, 10 minutes for extraction and purification, 5 minutes of pre-incubation of RNP T1, 5 minutes of pre-activation with sample, and a 10-minute dual-blocking reaction and readout step (Fig. 6a). Across biological replicates, our assay successfully detected MRSA in spiked blood samples down to 200 CFU per 200 µL (Fig.6b), yielding SNR values ranging from 1.3 to 12.9 for positive and 0.96 to 1.29 for negative value (Fig. 6c). Using receiver operating characteristic (ROC) curve, threshold SNR was established as 1.293, the assay correctly classified positive and negative samples with a 100 percent sensitivity and specificity, clearly separating true positives from true negatives.

**Figure 6.**
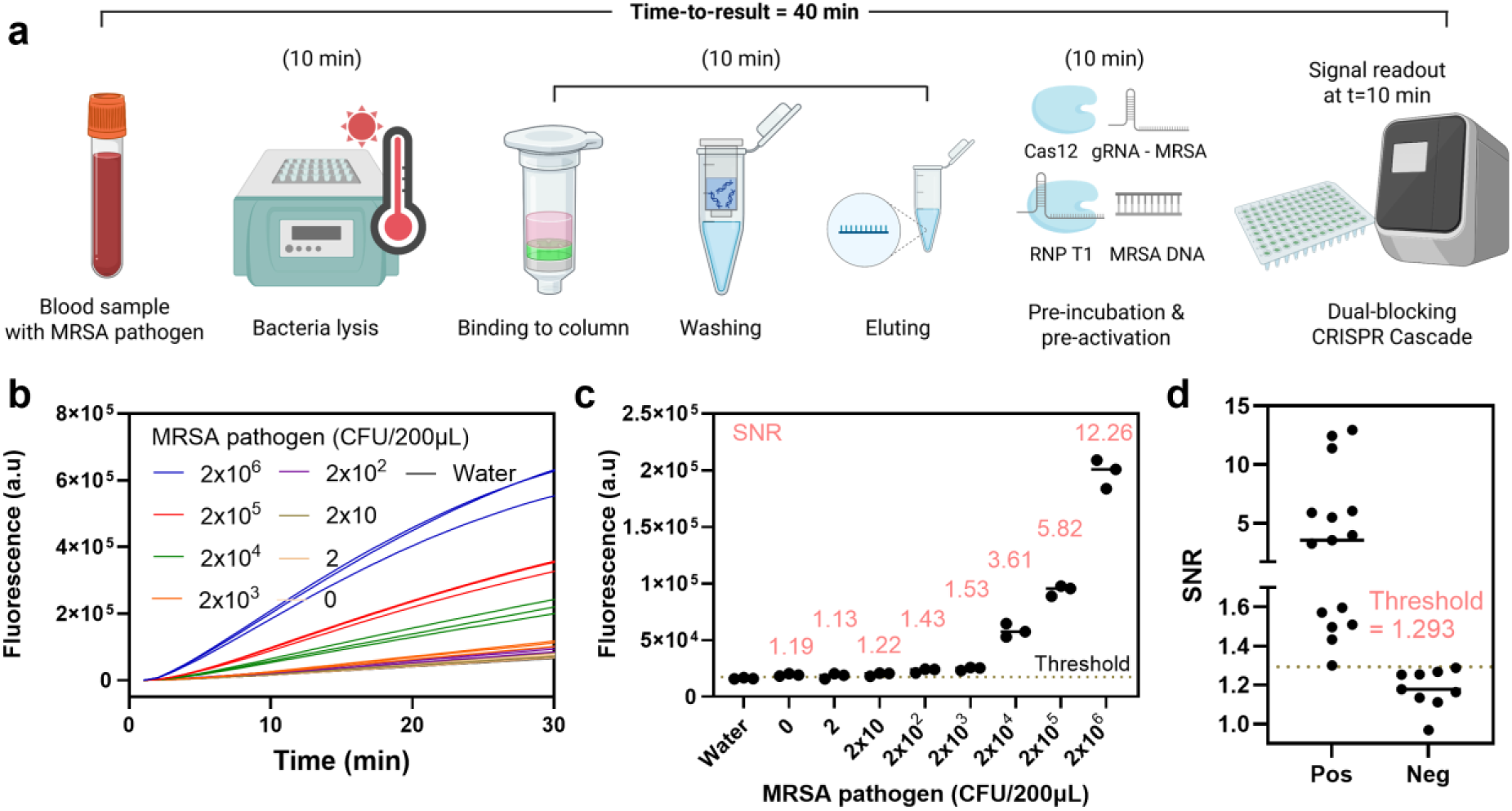
Evaluation of the dual-blocking CRISPR Cascade assay using whole-blood samples spiked with MRSA. (a) Workflow for mock clinical testing with MRSA-spiked whole blood, involving thermal lysis, column-based DNA extraction, and dual-blocking CRISPR Cascade detection, completed within 40 minutes. (b) Time-resolved fluorescence profiles of the dual-blocking CRISPR Cascade reactions over 30 minutes across varying MRSA concentrations, showing progressively increased signal intensity with higher pathogen loads. (c) End-point fluorescence and corresponding signal-to-noise ratio (SNR) values measured at 10 minutes, with SNR values highlighted in red. (d) Distribution of signal-to-noise ratio (SNR) values for positive and negative samples. Receiver operating characteristic (ROC) analysis established a threshold SNR of 1.293. Data is shown with individual well replicates (n = 3).

The observed sensitivity was limited by DNA loss during column-based purification, which reduces assay input efficiency at low pathogen concentrations. In particular, samples spiked with 2 and 20 CFU per 200 µL were classified as negative in the PCR analysis (Fig. S21), indicating that the reduced sensitivity primarily arose from extraction-related sample loss rather than from any limitation of the dual-blocking assay itself. Thus, coupling this autocatalytic CRISPR system with loss-free sample-handling strategies, such as our previously reported biphasic preparation protocol, may enable recovery of these ultra-low targets and further enhance real-world performance.^32–35^ Taken together, these findings show that the dual-blocking CRISPR-Cascade integrates robustly with routine clinical sample-processing methods, delivering rapid (∼40 min) and sensitive MRSA detection from purified whole-blood samples.

## Discussion

We demonstrate that dual-blocking both gRNA and trigger DNA provides two orthogonal kinetic gates, sharply reducing background activation (3–18-fold) while preserving atto-molar sensitivity. This strategy substantially reduces nonspecific activation in autocatalytic CRISPR reactions by imposing two sequential kinetic thresholds, thereby limiting spontaneous RNP formation and boosting signal discrimination (Fig. 4d). Our findings demonstrate that the dual-blocking strategy reduces background activation by approximately 3- to 18-fold compared to single-blocking approaches, while still preserving rapid signal generation and enabling sensitive, highly specific detection down to atto-molar levels. These findings expand the design principles for leakage-suppressed CRISPR circuits, highlighting the importance of coordinated control over both gRNA and trigger-DNA inputs in self-amplifying reactions.

This dual-blocking architecture functions as a molecular analogue of split-activator logic. Whereas traditional split activators typically divide a single activator substrate into two fragments that require co-localization,^36,37^ our design applies a similar principle across two orthogonal biochemical species: the guide RNA and trigger DNA. Looking forward, this architecture could be further generalized. For example, splitting the trigger DNA into two fragments, each masked by its own bubble-DNA, would produce a triple-blocking configuration. Such tri-layer gating could further enhance kinetic insulation that further suppresses unintended RNP formation. Systematically evaluating higher-order gating architecture represents an intriguing direction for minimizing nonspecific signals in autocatalytic systems.

Despite strong suppression in the dual-blocking configuration, residual leakage was still observed. This background likely arises from enzyme-driven strand invasion events: Cas12a can unwind scgRNA to form functional RNPs, and active RNPs can strand-invade Cascade probes,^17,18^ gradually exposing trigger sequences over time (Fig. S1). These leakage pathways appear difficult to eliminate solely through thermodynamic stabilization, suggesting that autocatalytic CRISPR circuits reach a biochemical ceiling in leakage suppression when relying only on nucleic-acid-based gates. Notably, introducing a large molar excess of sacrificial gRNA-T1 created a kinetic sink for Cas12a, diverting it away from blocked precursors and nearly abolishing background activation. This observation points to hybrid gating strategies combining dual nucleic-acid caging and saturating enzyme competition as a promising direction for approaching truly zero-leakage behavior. Looking ahead, it will be valuable to develop generalizable RNP T2 components paired with sequence-agnostic decoy guides that function independently of target identity. Strategies such as truncated guides or spacer-only RNAs lacking scaffold domains may offer a path toward universal decoy designs capable of suppressing leakage across diverse targets through improved nuclease specificity.^38,39^

Our architecture also streamlines point-of-care operation. Although a two-step configuration was used here to maximize performance, this design introduces a minor contamination risk similar to but less severe than conventional two-pot pre-amplification workflows. This limitation can be readily mitigated through strategies such as closed tube integration, physical barriers, or lyophilized reagents, which would enable a true one pot implementation without compromising sensitivity.^40^ To be specific, because Cas12a and gRNA can be pre-assembled into RNP T1 and lyophilized directly, the pre-incubation step can be eliminated entirely.^41–43^ RNP T2 precursors can likewise be pre-formulated, enabling a simple workflow in which both modules are rehydrated, mixed with the sample, and incubated isothermally. Future work could focus on optimizing lyophilization and stabilization chemistries to support robust storage, transport, and deployment in resource-limited settings.

An important consideration in scaling this technology is the increased design space associated with dual-blocking. Compared to single-blocking systems, dual-blocking introduces multiple tunable components, including the stoichiometry of trigger DNA to bubble DNA and ratios of RNP T1 to RNP T2 precursors. These variables offer opportunities to tailor dynamic range, kinetics, and specificity but also introduce optimization complexity.^28^ To systematically navigate this parameter space, reaction-kinetic modeling and computational design frameworks will be valuable.^44,45^ In-silico simulations that incorporate strand-exchange kinetics, R-loop formation, and enzyme–substrate competition could guide experimental design by predicting conditions that minimize leakage while maintaining amplification gain.^46,47^ Coupling such predictive modeling with targeted experimental screening will accelerate parameter identification and provide a rational foundation for optimizing multi-layer CRISPR autocatalytic circuits. However, current models are limited by incomplete understanding of the precise secondary structures that undergo strand invasion, making it challenging to fully capture unwinding dynamics in silico. Continued experimental characterization of blocked state conformations will therefore be critical for improving model accuracy and guiding next generation circuit design. Another significant limitation lies in accurately modeling RNA–DNA duplex formation.^48–50^ Current computational frameworks are primarily optimized for DNA–DNA or RNA–RNA interactions, and the hybridization thermodynamics and secondary structure prediction for RNA–DNA hybrids remain poorly parameterized. The lack of experimentally validated energy models and kinetic constants for such heteroduplexes makes in-silico integration of RNA strands challenging. Developing improved hybrid thermodynamic models will be crucial for advancing predictive design of mixed nucleic acid systems in CRISPR autocatalytic circuits. Ultimately, integrating computational and empirical approaches will be essential for refining next-generation self-amplifying CRISPR platforms and expanding their translational readiness.

Overall, dual-blocking establishes a generalizable blueprint for constructing self-amplifying CRISPR systems with PCR-like binary discrimination, yet without thermal cycling. By combining orthogonal gating, sacrificial competitor strategies, and systematic optimization, our approach addresses fundamental leakage challenges that have limited previous self-amplifying CRISPR assays. Our design improves practical specificity by maintaining a completely flat baseline in the absence of target, thereby enabling clear on–off switching behavior.

## Conclusion

This work establishes a dual-blocking autocatalytic CRISPR-Cascade architecture that addresses the intrinsic leakage mechanisms that have limited previous amplification-free CRISPR systems. By independently caging the guide RNA and the trigger DNA, the system introduces two orthogonal kinetic barriers that markedly suppress spontaneous RNP formation, while preserving rapid detection and atto-molar sensitivity. Incorporation of a sacrificial guide RNA decoy adds a third AND gate component by acting as a kinetic sink for Cas12a, further reducing unintended activation and driving the baseline toward PCR-like flat behavior in the absence of target. Together, these multilayer strategies enable robust MRSA pathogen detection from purified whole blood within 40 minutes and achieve complete classification accuracy in spiked samples, demonstrating practical diagnostic utility. More broadly, the modularity and generalizability of the dual-blocking design provide a framework for constructing next generation self-amplifying CRISPR circuits with reduced background noise and reliable AND gate switching suitable for diverse applications in molecular sensing and point of care diagnostics.

## Materials and Methods

### Reagents

PBS buffer (21040CV) was obtained from Corning. 10x TBE buffer (B52), UltraPure™ 1 M Tris-HCI Buffer (15567027)-pH 7.5, 5 M NaCl (2802039), and RNase-free water (AM9760G) were purchased from Thermofisher Scientific. 1X TE-pH 8.0, NEB low molecular weight DNA ladder (N3233S), Magnesium Chloride (MgCl2, B9021S), Murine RNase Inhibitor (M0314L), and NEBuffer™ r2.1 (5 mL, B6002S) solution were purchased from New England Biolabs (NEB). Alt-R™ LbCas12a (Cpf1) Ultra and all nucleic acid sequences were synthesized and obtained from Integrated DNA Technologies (IDT).

### Pathogens and DNA sequences

The synthetic MRSA DNA fragment was synthesized by IDT and dissolved in TE buffer. Genomic DNA from the MRSA strain HFH-30106 (NR-10320) was procured from BEI Resources (Manassas, VA), aliquoted to the required concentrations and volumes, and stored at −80 °C until use. For bacterial culture experiments, MRSA strain HFH-30106 (NR-10192) was likewise obtained from BEI Resources.

### Switchable caged guide RNA and Cascade probe design and sequences

The DNA and RNA strands listed in SI Appendix, Table S1 were shipped in lyophilized form and resuspended in Tris-EDTA (TE) buffer. 1µM Cascade probes (CP) were prepared by hybridization the two complementary oligomers at the molar ratio of 1:1.5 of (trigger DNA:bubble DNA) at 95°C for 4 min in annealing buffer [20 mM tris-HCl (pH 7.5), 50 mM NaCl, and 10 mM MgCl2], followed by gradient cooling to 25°C at a rate of 0.1°C/s. 1 µM switchable caged guide RNA (scgRNA) solutions were also generated by the hybridization of the gRNA (gRNA T2), inhibitor-handle (iH) DNA, and inhibitor-spacer (iS) DNA using the same annealing procedure with the molar ratio of 1:1.2:1.2. The concentration of CP and scgRNA specified in the text refers to the concentration of the used gRNA and TD. All scgRNAs and CPs were prepared, aliquoted, and stored at −20 °C until use. Secondary-structure predictions for BNAs (i.e., CP and scgRNA constructs) were performed using NUPACK software under ionic conditions of 50 mM Na⁺ and 10 mM Mg²⁺ at 37°C. Because NUPACK does not support hybrid secondary-structure prediction between RNA and DNA strands within a single complex, the RNA guide sequences (gRNA) were converted to their DNA equivalents prior to simulation. This substitution preserves the base-pairing pattern and thermodynamic characteristics of the nucleic acid structures, allowing approximate estimation of the hybridization behavior and folding energy landscapes of the designed scgRNA complexes. For melting curve analysis, 1x Evagreen (Biotium) was added to each sample with the ratio of 9:1 (Evagreen:sample). Then, the melting curve was recorded using a QuantStudio3 (Applied Biosystems) instrument with a controlled temperature ramp rate of 0.1°C/s.

### Single CRISPR Cas12a assay

The following protocol was applied to all single-Cas12a experiments, including the 2×2 matrix characterization shown in Figure 2f and Figures 3c–3e. In this format, each reaction consisted of one Cas12a enzyme complexed with a corresponding guide RNA and the target nucleic acid, which could be either in blocked or unblocked form. Specifically, the i-a condition in Figure 2f represents the reaction containing gRNA-T2 and trigger DNA, while the i-b condition represents gRNA-T2 with the Cascade probe, where the target DNA is replaced with its blocked form. The ii-a condition corresponds to scgRNA with trigger DNA, and the ii-b condition to scgRNA with the Cascade probe, where both the gRNA and target are in blocked states. All single-Cas12a reactions were pre-incubated for 20 min at 37°C with 10 nM LbCas12a, 10 nM gRNA, 0.4 U µL⁻¹ Murine RNase Inhibitor, 500 nM fluorescent ssDNA reporter, 1×NEB buffer, and 2.5 mM MgCl₂. After pre-incubation, 9 µL of the reaction mixture was combined with 2 µL of the sample containing varying concentrations of target nucleic acids. The reaction was conducted on a QuantStudio3 real-time fluorescence reader, and fluorescence intensities were recorded every minute. Fluorescence data were normalized by subtracting the initial (0 min) fluorescence signal, and the signal-to-noise ratio was calculated by dividing the fluorescence signal of the positive group by that of the negative control at the time point of interest, in which RNase-free water served as the sample. The same conditions were used for RNP T2-only characterization experiments, except that RNP T1 was replaced by RNP T2 and the target molecules were substituted with the trigger DNA or Cascade probe.

### High-resolution non-denaturing polyacrylamide gel electrophoresis (PAGE) assays

High-resolution non-denaturing polyacrylamide gel electrophoresis (PAGE) experiments were prepared with 15% polyacrylamide in 10xTBE (89 mM Tris, 89 mM boric acid, 2 mM EDTA). Gel solution was prepared by mixing 3.75 mL Biorad 40% Acrylamide/Bis Solution, 5.25 mL degassed water, 1 mL 10X TBE (ThermoFisher, 161-0733), 50 μL APS (1610700, 10% w/v solution), and 5 μL TEMED (1610800). NEB Low Molecular Weight Ladder (N3233S) was used as the DNA standard. The trans-cleavage activity of RNP T1 on scgRNA and Cascade probes (CP) was evaluated using non-denaturing polyacrylamide gel electrophoresis (PAGE). Cleavage reactions were quenched by transferring to ice, adding 2 µL of loading dye (with a ratio of 8 µl of RNP T1 solution to 2 µl of loading dye) and immediately loaded into the designated wells of the 15% acrylamide gel. Electrophoresis was carried out at room temperature under a constant voltage of 130 V for 3-4 h. Following electrophoresis, gels were stained with ThermoFisher SYBR 1 gel stain (S7563) prepared at a ratio of 5 µL 10×SYBR to 50 µL 1×TBE buffer, and gently agitated at 150 rpm in the dark for 40 min. Gels were imaged using a Bio-Rad gel documentation system.

### CRISPR-Cascade assay

For the dual-blocking Cascade assay, the RNP T1 complex, consisting of 10 nM LbCas12a, 10 nM target specific gRNA, 2.5 mM MgCl₂, and 1×NEB 2.1 buffer, was preincubated at 37 °C for 5 minutes to allow RNP formation. The preassembled RNP T1 was then activated by mixing with pathogen genomic DNA at 37°C for 5 min in a 1:1 volume ratio of RNP T1 to sample. Separately, RNP T2 complexes were prepared with 10 nM LbCas12a, 0.4 U/µL Murine RNase Inhibitor, 500 nM DNA fluorescent probe, 1× NEBuffer™ r2.1, and an additional 2.5 mM MgCl₂, and stored at 4°C until use. After activation of RNP T1, RNP T2 was supplemented with 10 nM scgRNA and 10 nM Cascade probe (CP) to complete the reaction mixture. The completed RNP T2 complex was then combined with the activated RNP T1 at a 3:1 volume ratio.

For assay development and performance characterization, the concentrations of RNP T1 and RNP T2 were kept constant, while their mixing ratio was systematically varied from 1:1 to 1:48. Reactions were performed on a QuantStudio3 instrument with fluorescence signals recorded at 1-minute intervals. Fluorescence readings were baseline-corrected by subtracting the initial (0-minute) value, and the signal-to-noise ratio was determined by dividing the fluorescence intensity of the positive sample by that of the negative control. For each single blocking Cascade assay in Figs. 4c and 4d, all groups were tested under identical reaction conditions, with the only difference being that one blocking component (either the scgRNA or Cascade probe) was substituted by its unblocked form (gRNA or trigger DNA) at the same concentration to allow direct comparison. For the gRNA decoy and multi-guide enhancement strategies, additional gRNA-T1 or gRNA-T2-310 (Supplementary Table 1) was added to the T1 reaction tube, with an equal volume removed from the water component to maintain a consistent total reaction volume and overall reagent concentrations.

### Pathogen culture

Tryptic soy broth (TSB) and agar were obtained from the Cell Media Facility at the University of Illinois Urbana–Champaign (UIUC). Methicillin-resistant *Staphylococcus aureus* (MRSA) was cultured in TSB and incubated at 37 °C for 16 h. After incubation, phosphate-buffered saline (PBS) stocks were prepared by centrifuging 1 mL of the overnight culture at 6,000 × *g* for 10 min to pellet the cells. The pellet was washed twice with 1×PBS, resuspended in 1 mL PBS, aliquoted, and stored at room temperature. Stocks were used within four days of preparation. Colony counts were verified by plating, and appropriate dilutions were made in whole blood for experimental use.

### Detection of Pathogen in Blood Sample

Venous whole blood was obtained from BIOIVT (HUMANWBK2-0101184; Human Whole Blood K₂EDTA, gender unspecified) and stored at 4 °C on a rotating mixer for up to two weeks before experimentation. To achieve target bacterial concentrations, tenfold serial dilutions of bacterial stocks were prepared and spiked into whole blood. Genomic DNA was extracted using the QIAamp DNA Blood Kit (QIAGEN, 51104) following the manufacturer’s protocol. Briefly, 200 µL of blood was treated with QIAGEN Protease and incubated with lysis buffer at 56 °C for 10 min. Ethanol was added to the lysate, which was then transferred to a QIAamp Mini spin column. After centrifugation, DNA bound to the column was washed sequentially with kit-provided buffers to remove impurities and subsequently eluted in 200 µL nuclease-free water by centrifugation. The sample was measured with the CRISPR-Cascade assay.

### Polymerase chain reaction (PCR)

PowerTrack SYBR Green Master Mix (Thermo Fisher Scientific) was prepared following the manufacturer’s protocol. qPCR reactions were conducted in a 10 μL total volume containing 1 μL of sample and 200 nM each of forward and reverse primers. A standard calibration curve was generated using MRSA genomic DNA ranging from 1 to 10⁴ copies per μL in 10-fold serial dilutions. Amplification was performed on a QuantStudio3 Real-Time PCR System. Extracted DNA from whole-blood samples spiked with MRSA were classified as positive or negative based on whether their Ct values fell within the range defined by the standard calibration curve.

## Author Contribution

J.L., A.B.V., and M.W. equally contributed to this paper. J.L., E.V., and R.B., designed research; J.L., A.B.V., and M.W., K.K. performed the experiments; J.L., A.B.V., M.W., and R.B. analyzed data; and J.L., A.B.V., M.W., K.K., E.V., and R.B. wrote the paper.

## Supporting Information Available

This article contains supporting information online.

## Use of Artificial Intelligence

The initial draft of this manuscript was written by the authors. ChatGPT (OpenAI GPT 5, 2025 version) was used only to improve sentence clarity, grammar, and readability. All AI assisted edits were carefully reviewed and verified by the authors to ensure scientific accuracy and integrity.

## Acknowledgement

1. J. L. acknowledge Material Research Lab (MRL, UIUC) for the Professor Joe Greene Postdoctoral Fellowship by Material Research Laboratory at UIUC. The authors acknowledge BioRender for figure creation. Figures 1, 2e, 3b, 4a, 5a, and 6a were generated using BioRender. The authors thank the staff at the Holonyak Micro and Nanotechnology Laboratory at UIUC for facilitating the research and the funding from University of Illinois. The following product was obtained through BEI Resources, NIAID, NIH: NR-10320 and NR-10192. This work was partially supported by VinUni-Illinois Smart Health Center and the NIH (R01AI148385 & R01EB032725 A). R.B. and E.V. acknowledge partial support from the Jump ARCHES (Applied Research through Community Health through Engineering and Simulation) endowment through the Health Care Engineering Systems Center at UIUC and OSF.

