## Supporting Information for "Amplification-Free Dual-Blocking Autocatalytic CRISPR-Cascade for Atto-Molar DNA Detection with Low Nonspecific Signal"

**Supplementary Table 1.** Oligo nucleic acid sequences used in this paper.

| **Sequence Name** | **Sequence** | **Used in Figures** |
| --- | --- | --- |
| gRNA T2 | rUrArArUrUrUrCrUrArCrUrArArGrUrGrUrArGrArUrGrArUrCrGrUrUrArCrGrCrUrAr rArCrUrArUrGrA | 2,3,4,5,6 |
| iDHA – Handle | ATCTACACTT **TTTATTT** AGTAGAAATTA | 2,3,4,5,6 |
| iDNA – Spacer | TCATAGTTAG **TTTATTT** CGTAACGATC | 2,3,4,5,6 |
| Trigger DNA | TCATAGTTAGCGTCATAGTTAGCGTAACGATCTAAAGTTTT | 2,3,4,5,6 |
| Bubble DNA 41nt | AAAACTTTAGATCGTTACGCT **TTTATTT** AACTATGACGCTAACTATGA | 2,3,4 |
| Bubble DNA 35nt | TTAGATCGTTACGCTAAC **TTTATTT**  TATGACGCTAACTATGA | 3 |
| Bubble DNA 29nt | AAAACTTTAGATCGT **TTTATTT**  TACGCTAACTATGA | 3 |
| Bubble DNA 23 nt | TTAGATCGTTAC **TTTATTT**  GCTAACTATGA | 3 |
| 2 bubble-  Bubble DNA | AAAACTTTAGATCG **TTTATTT**  TTACGCTAACTATG **TTTATTT**  ACGCTAACTATGA | 3 |
| 3 bubble-  Bubble DNA | AAAACTTTAGA **TTTATTT**  TCGTTACGCT **TTTATTT**  AACTATGACG **TTTATTT**  CTAACTATGA | 3 |
| Bubble DNA  with 1 LNA | AAAACTTTAGATCGTTACGC +T **TTTATTT** +A  ACTATGACGCTAACTATGA | 3 |
| Bubble DNA  with 2 LNA | AAAACTTTAGATCGTTACG  +C +T **TTTATTT** +A +A  CTATGACGCTAACTATGA | 3 |
| Two parts Bubble DNA - Part A | AAAACTTTAGA **TTTATTT**  TCGTTACGCT | 3 |
| Two parts Bubble DNA - Part B | AACTATGACG **TTTATTT**  CTAACTATGA | 3 |
| New Trigger DNA | GAAACTCATAGTTAGCGTAACGATCTAAAG | 3 |
| New Bubble DNA | CTTTAGATCGTTACG **ATATA** CTAACTATGAGTTTC | 3 |
| gRNA T1  MRSA 1 | rUrArArUrUrUrCrUrArCrUrArArGrUrGrUrArGrArUrGrGrUrCrUrArArArArUrUrUrUrArCrCrArCrGrU | 5,6 |
| gRNA T1  MRSA 2 | rUrArArUrUrUrCrUrArCrUrArArGrUrGrUrArGrArUrUrArGrArUrCrUrUrArUrGrCrArArArCrUrUrArA | 5,6 |
| gRNA T1  MRSA 3 | rUrArArUrUrUrCrUrArCrUrArArGrUrGrUrArGrArUrUrCrUrUrUrArUrCrArUrArUrGrArUrArUrArArA | 5,6 |
| gRNA T1  MRSA 4 | rUrArArUrUrUrCrUrArCrUrArArGrUrGrUrArGrArUrUrArArUrGrCrGrCrUrArUrArGrArUrUrGrArArA | 5,6 |
| gRNA T2 310 | rUrArArUrUrUrCrUrArCrUrArArGrUrGrUrArGrArUrGrGrArUrArUrArCrGrArUrArUrArUrArUrArUrArU | S18 |
| MRSA gene Fragment | ACGTGGTAAAATTTTAGACCATCTACACTTAGTAGAAATTACCCTATAGTGAGTCGTATTA | S13 |
| Reporter | /56-FAM/TT TTT T/3IABkFQ/ | All |
| MRSA PCR  Primer F | GTAGAAATGACTGAACGTCCGATAA | S21 |
| MRSA PCR  Primer R | CCAATTCCACATTGTTTCGGTCTAA | S21 |

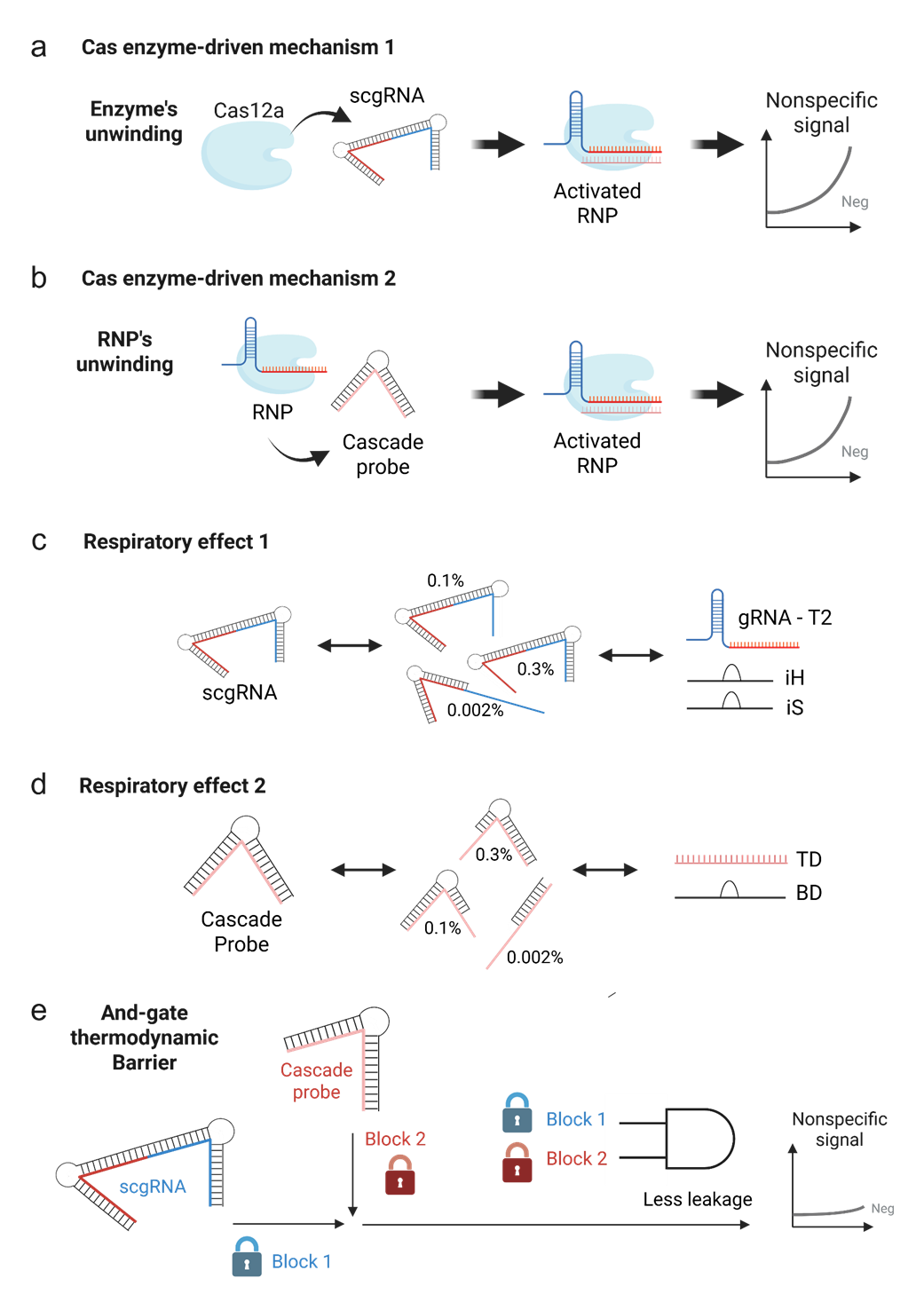

**Figure S1.** Leakage pathways in single-blocking systems and how dual-blocking suppresses nonspecific activation. (a) Cas12a can invade and partially unwind scgRNA to form functional RNPs that subsequently become activated upon encountering trigger DNA, generating background signal. (b) Pre-formed RNP T2 complexes can strand-invade and unwind Cascade probes, misrecognizing them as trigger DNA and initiating the autocatalytic feedback loop. (c-d) Additionally, both scgRNA and Cascade probes undergo thermodynamic breathing (i.e., respiratory effect), transiently sampling partially unblocked conformations that seed spontaneous activation. (e) To mitigate these leakage mechanisms, the dual-blocking architecture simultaneously cages both gRNA and trigger DNA using scgRNA and Cascade probes, requiring cleavage of both bulge elements prior to productive RNP T2 assembly. In other words, even if one component transiently adopts an active conformation due to stochastic structural fluctuations, the system remains inactive as long as the other component stays blocked. This interdependency prevents unintended activation, ensuring that nonspecific amplification cannot proceed unless both molecular gates are simultaneously satisfied. This AND-gate logic imposes two orthogonal thermodynamic and kinetic barriers to spontaneous activation, markedly reducing leakage and nonspecific background relative to single-blocking systems.

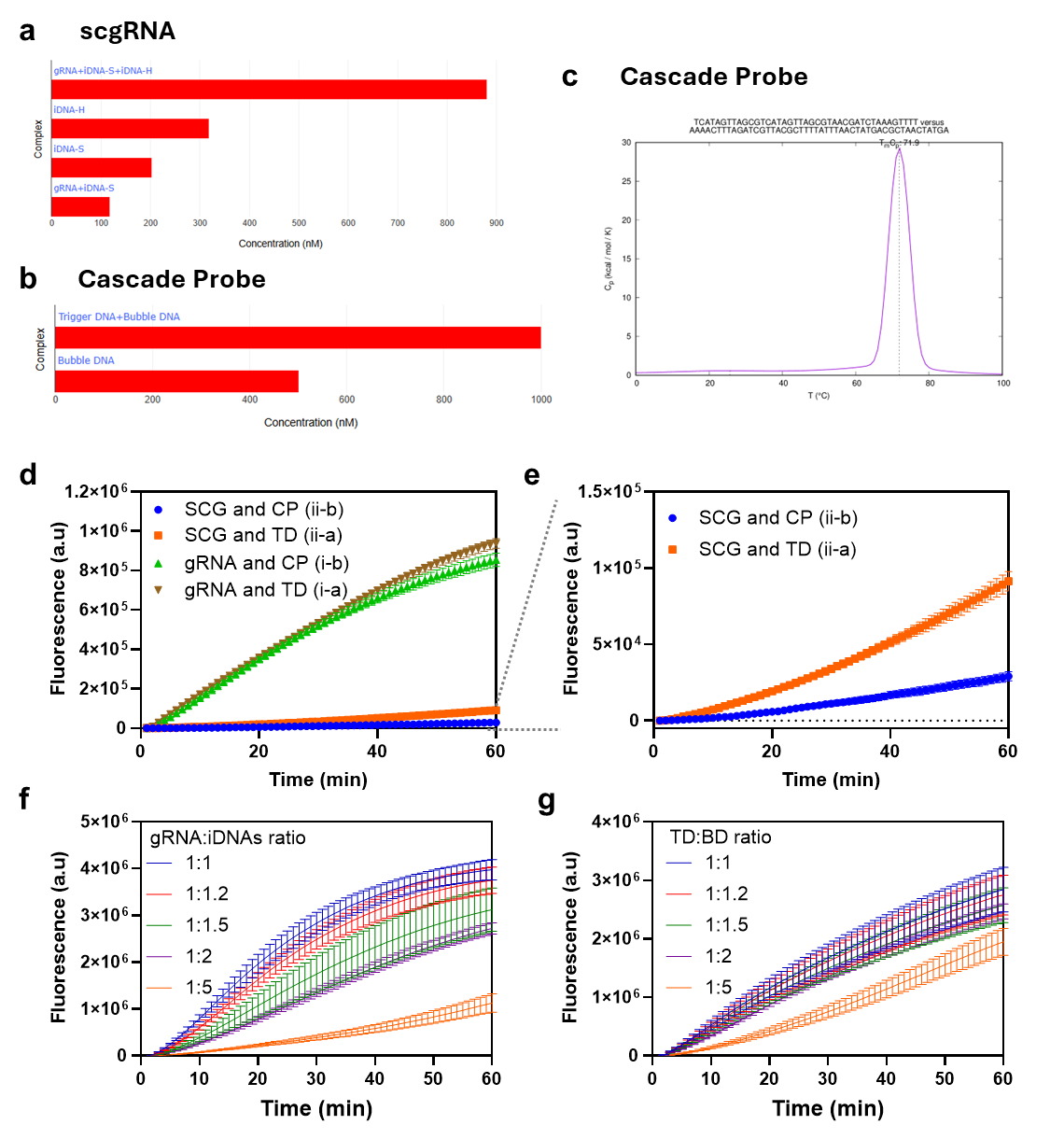

**Figure S2.** **Thermodynamic validation and blocking behavior of scgRNA and Cascade probe designs.** (a–b) NUPACK-predicted duplex formation efficiencies for scgRNA and Cascade probe components under Na⁺ (50 mM) and Mg²⁺ (10 mM) conditions. scgRNA was formed using 1 µM gRNA-T2 with 1.2 µM each of iDNA-handle (IH) and iDNA-spacer (IS), and the Cascade probe was formed using 1 µM trigger DNA (TD) and 1.5 µM bubble DNA (BD). For modeling purposes, gRNA-T2 sequence was represented as DNA to enable base-pairing simulation. (c) DINAMelt prediction of Cascade probe melting temperature (~71.9°C), consistent with experimental melting curves in Fig. 2c. (d) Raw fluorescence traces and (e) expanded views of the 2×2 single-Cas12a assay (matching Fig. 2f), isolating RNP T2 formation under four blocking states (unblocked, TD-blocked, gRNA-blocked, dual-blocked). Dual-blocking yielded the lowest signal trajectory, confirming minimal spontaneous activation. (f–g) Increasing blocker ratios for scgRNA (gRNA:iDNAs) and Cascade probe (TD:BD) from 1:1 up to 1:2 reduced leakage but did not completely eliminate background activation, indicating that leakage does not arise solely from insufficient blockade strength. Even though a 1:5 ratio further suppressed the noise, this suppression arises not only from shifting the reaction toward a strongly blocked equilibrium state but more dominantly from the excess iDNAs or BD molecules, which act as competitive inhibitors by nonproductively sequestering RNPs and thereby attenuating their catalytic turnover in trans-cleavage reactions. These results support that Cas12a-driven unwinding constitutes a kinetic bypass mechanism rather than a purely thermodynamic deficiency, reinforcing the need for a dual-blocking architecture to attenuate seed formation that triggers autocatalytic reaction.

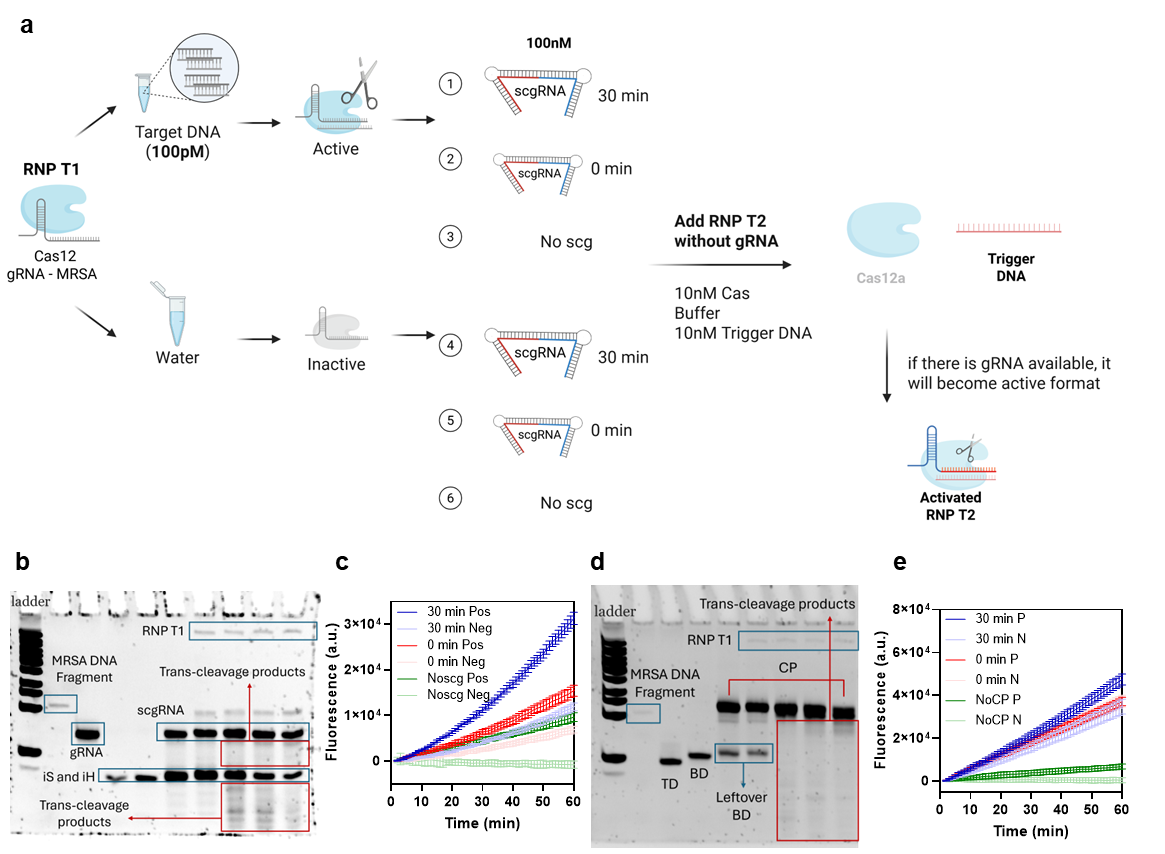

**Figure S3.** Activated RNP T1 cleaves scgRNA and Cascade probes to release functional nucleic acids that initiate downstream autocatalysis. (a) Workflow used to examine whether activated RNP T1 can cleave single-stranded DNA bulges within scgRNA and Cascade probes, releasing functional nucleic acids capable of initiating downstream autocatalysis. RNP T1 was pre-incubated with either 100 pM MRSA DNA (activated) or water (inactive control), then mixed with scgRNA or Cascade probe for 0 or 30 min, or omitted entirely (no-scg negative control). Cleavage of the bulge regions was expected to liberate gRNA or trigger DNA that could assemble with Cas12a to form new active RNPs (RNP T1 + newly formed RNP T2), resulting in enhanced fluorescence signal in the reporter assay. (b) Non-denaturing PAGE verified time-dependent trans-cleavage of scgRNA by activated RNPT1. The scgRNA was generated by hybridizing 1 μM gRNA with 1.2 μM iHandle (iH) and 1.2 μM iSpacer (iS) to yield 1 μM scgRNA, then diluted to 100 nM. The RNPT1 complex was formed by incubating 10 nM Cas12a with 10 nM gRNA-MRSA for 20 min at 37°C, followed by activation with 20 nM MRSA DNA for 10 min. 9 µl of activated RNPT1 was subsequently incubated with 1 µl of 1 µM scgRNA at 37°C, and reactions were stopped at 0, 30, 60, and 120 min. Lanes 1–6 contained 1) the NEB small molecular ladder, 2) MRSA DNA fragment (20 nM), 3) gRNA (100 nM), 4) iSpacer (100 nM), 5) iHandle (100 nM), and 6) scgRNA (100 nM), respectively. Lane 6 contained scgRNA and unreacted iS/iH. Lane 7 included RNP T1, MRSA DNA, and scgRNA at 0 min. Lanes 8–10 represented trans-cleavage reactions at 30, 60, and 120 min. At 30 min (lane 8), partial scgRNA cleavage produced smaller DNA fragments. As incubation proceeded (lanes 9–10), progressive degradation of scgRNA, iS, and iH was observed, accompanied by accumulation of low–molecular-weight fragments. The time-dependent loss of scgRNA and increase in short species confirmed continuous RNPT1 trans-cleavage activity. Released gRNA fragments were detected alongside degradation of free and released iS/iH, demonstrating RNP T1’s collateral nonspecific cleavage activity, which likely competed with intact scgRNA over time. (c) Fluorescence measurements supported the gel data, showing an increase in signal when activated RNP T1 was incubated with scgRNA, peaking at 30 min, consistent with time-dependent liberation of functional gRNA. The 0-min condition displayed an intermediate signal, reflecting partial cleavage upon immediate mixing, whereas the no-scg control exhibited baseline fluorescence corresponding to RNP T1 alone. (d) Non-denaturing PAGE analysis of Cascade probe (CP) cleavage showed a parallel trend. CP was prepared by hybridizing 1 μM Trigger DNA (TD41) with 1.5 μM Bubble DNA (BD41) to generate 1 μM CP, then diluted to 100 nM. RNP T1 assembly and activation followed the same conditions as for scgRNA. 9 µl of activated RNP T1 was incubated with 1 μl of 1 μM CP at 37°C, and reactions were stopped at 0, 30, 60, and 120 min. Lanes 1–4 contained the 1) NEB small molecular ladder, 2) MRSA DNA fragment (20 nM), 3) TD (100 nM), and 4) BD (100 nM), respectively. Lane 5 included CP and unreacted TD/BD, while lane 6 represented RN PT1, CP, and unreacted components at 0 min. Lanes 7–9 corresponded to 30, 60, and 120 min trans-cleavage reactions. Partial CP cleavage was observed at 30 min, generating short fragments; extended incubation resulted in more degradation of CP, TD, and BD, accompanied by accumulation of low–molecular-weight DNA fragments. The TD and BD band gradually disappeared with time, indicating total RNPT1-mediated degradation. (e) Corresponding fluorescence measurements with Cascade probe, showing the same trend as scgRNA: activated RNP T1 incubation increased signal in a time-dependent manner, confirming that trigger DNA released from Cascade probes can form active RNP T2 complexes.

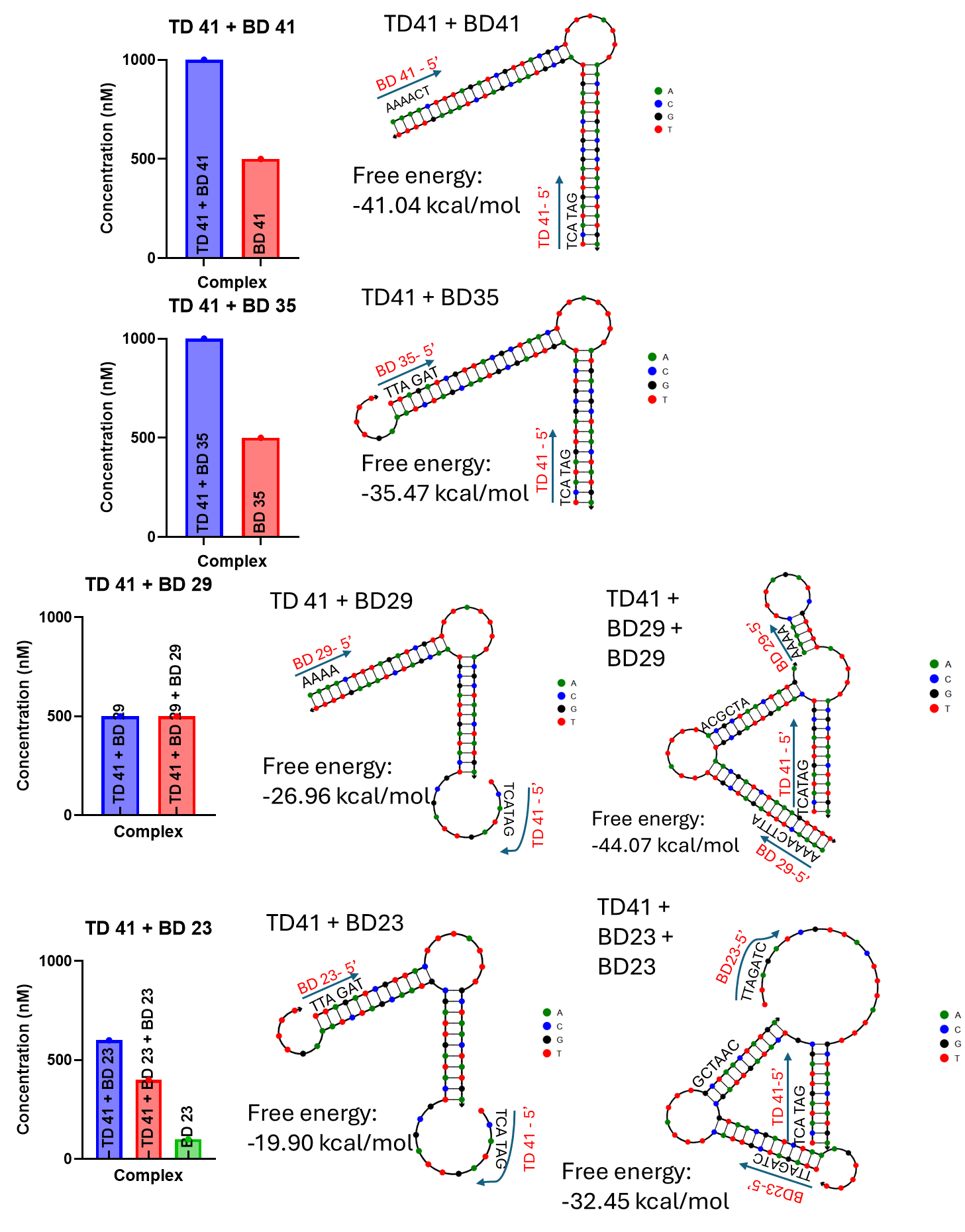

**Figure S4.** NUPACK-predicted secondary structures of Cascade-probe variants with progressively shortened bubble-DNA scaffolds. Each row shows results obtained when 1 µM trigger DNA (TD) and 1.5 µM bubble DNA (BD) were hybridized under 50 mM Na⁺ and 10 mM Mg²⁺ at 37°C. The left panels display the resulting duplex compositions, while the right panels present NUPACK-predicted secondary structures with annotated 5′ termini and corresponding free-energy values. When multiple stable duplexes were predicted, all are shown. Cascade probes were engineered by truncating the flanking duplex regions while preserving the central 7-nt bulge motif (TTTATTT). TD was fixed at 41 nt, while BD length decreased progressively from 41 nt to 35, 29, and 23 nt. BD35 retained the bulge sequence from BD41 but was shortened by 3 nt at both the 5′ and 3′ ends (total –6 nt), and subsequent variants (BD29 and BD23) were further truncated by 6 nt each. The 5′ ends and representative nucleotides of TD and BD are labeled for clarity.

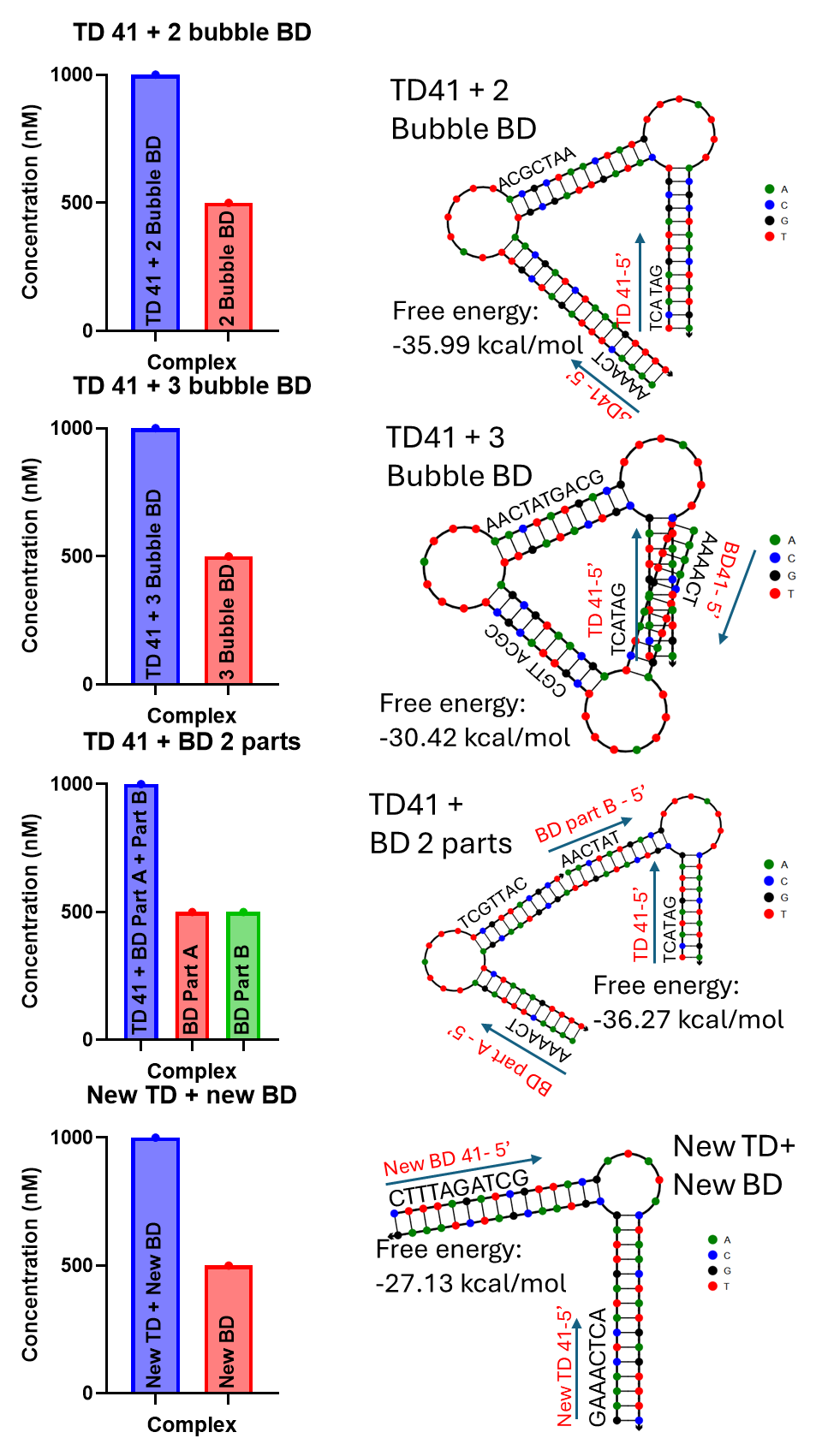

**Figure S5.** NUPACK-predicted structures and functional screening of Cascade-probe variants with altered bulge number, duplex segmentation, and new TD and BD sequence. LNA-modified probes, not included in the figure, (LNA-1 and LNA-2) are same structure with TD41 + BD41 in Figure S4, except that there are one or two LNA at the feet of the bulge.

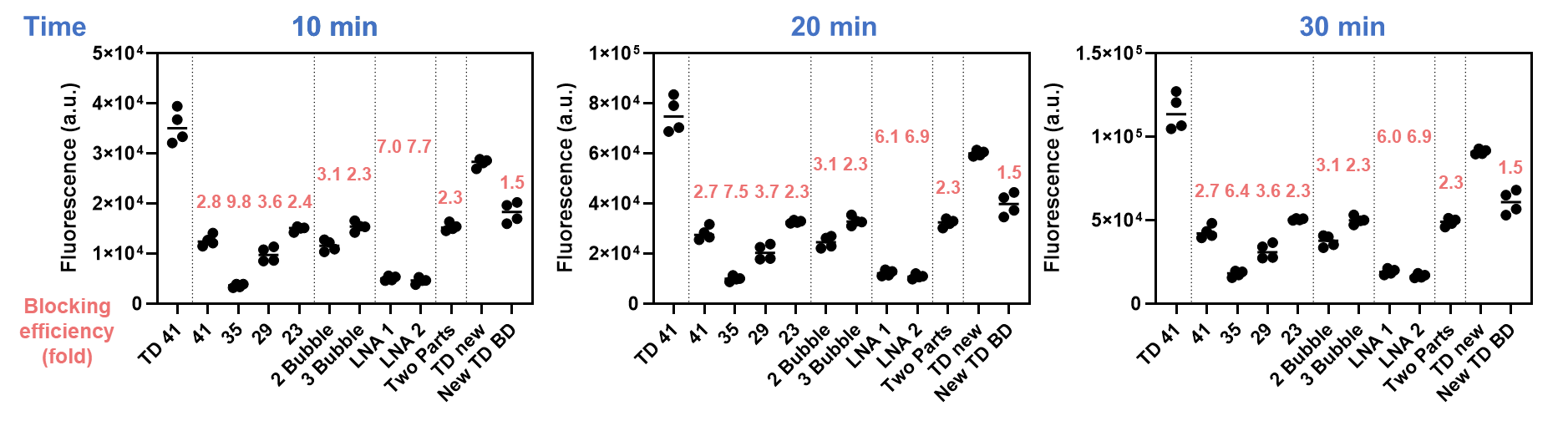

**Figure S6.** End-point fluorescence values for Cascade probe variants at 10, 20, and 30 minutes in the single-Cas12a assay. Bubble-DNA lengths of 41, 35, 29, and 23 nt represent designs shortened in 6-nt increments (excluding bulges); “2-bubble” and “3-bubble” indicate the number of bulge units; “LNA-1” and “LNA-2” denote incorporation of one or two LNAs beneath the bulge; and “two-parts” indicates division of the bubble-DNA into two segments. Blocking efficiencies, calculated as the fluorescence ratio relative to the unblocked TD41 condition, are displayed in red for each time point and each variant. Data are shown with individual replicates (n = 4).

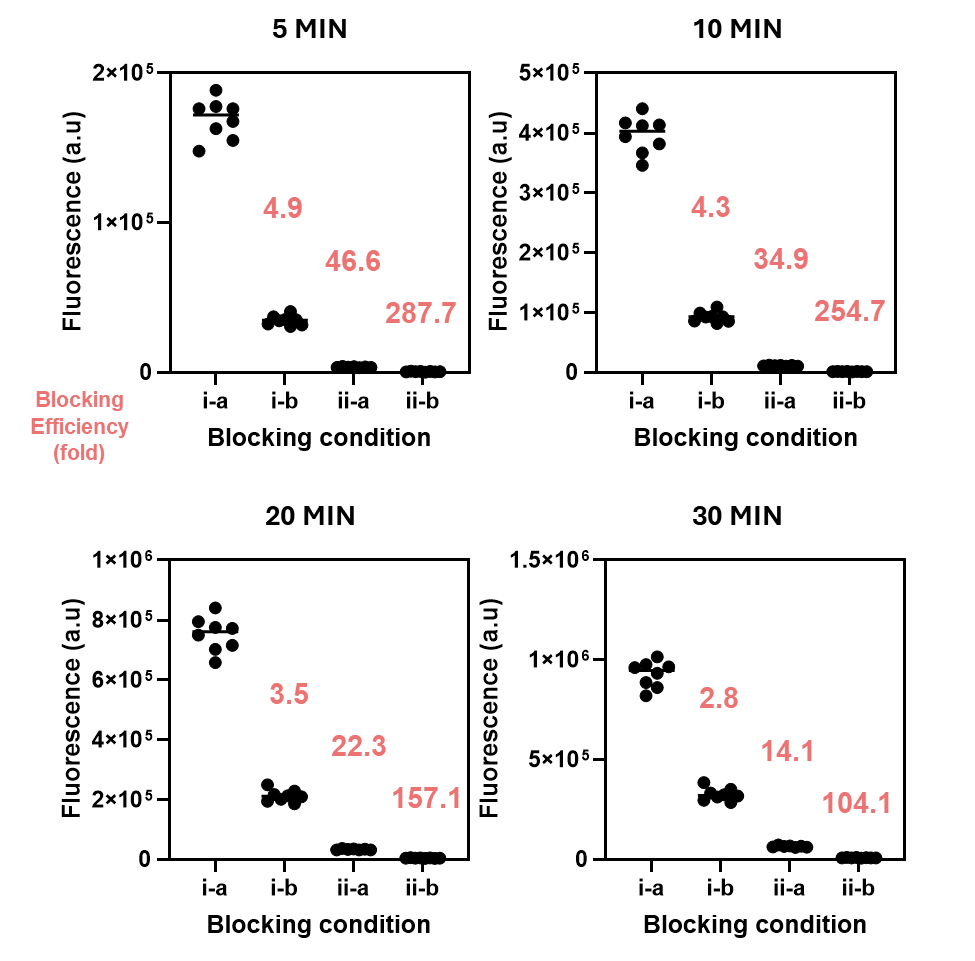

**Figure S7.** Blocking efficiency summary for the four precursor states in the single-Cas12a assay using the LNA-1 Cascade probe variant (unblocked, Cascade-blocked, scgRNA-blocked, and dual-blocked). Blocking efficiencies (shown in red) are reported for each condition at 5, 10, 20, and 30 min. Dual-blocking displayed the greatest suppression across all time points, followed by scgRNA-blocking and Cascade-blocking. Data are shown with individual replicates (n = 8).

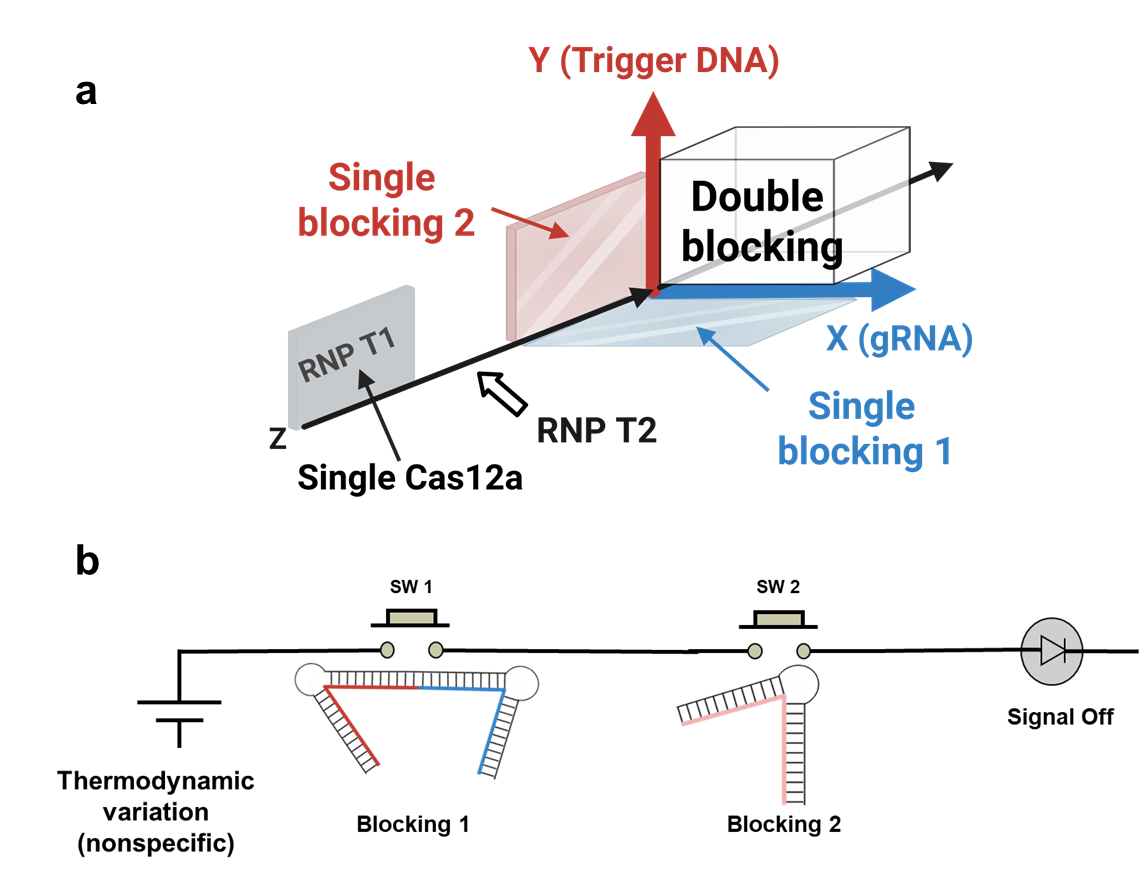

**Figure S8.** Sequential AND-gate mechanism underlying dual-blocking architecture. (a) Conceptual circuit diagram comparing single-blocking and dual-blocking configurations. Activated RNP T1 feeds into the RNP T2 autocatalytic module. Blocking can be applied on the gRNA axis (single blocking 1) or on the trigger-DNA axis (single blocking 2), and both can be applied simultaneously at their intersection to implement dual-blocking. This configuration functions as a sequential AND-gate, where both inhibitory modules must be released to fully activate the autocatalytic loop. The combined action of the two blocking pathways exhibits measurable synergistic suppression, exceeding the expected product of their individual-blocking efficiencies and reinforcing the cooperative behavior observed in Fig. 3. (b) Schematic illustration showing that dual-blocking operates as a sequential AND gate, where scgRNA sequestration functions as the primary kinetic checkpoint followed by Cascade-probe activation. Because Cas12a must first bind gRNA to form an active RNP before engaging trigger DNA, gRNA blocking imposes the dominant barrier to nonspecific activation. As a result, the system behaves as a serial switch, preventing leakage even under thermodynamic fluctuations and despite improvements in trigger-blocking stability.

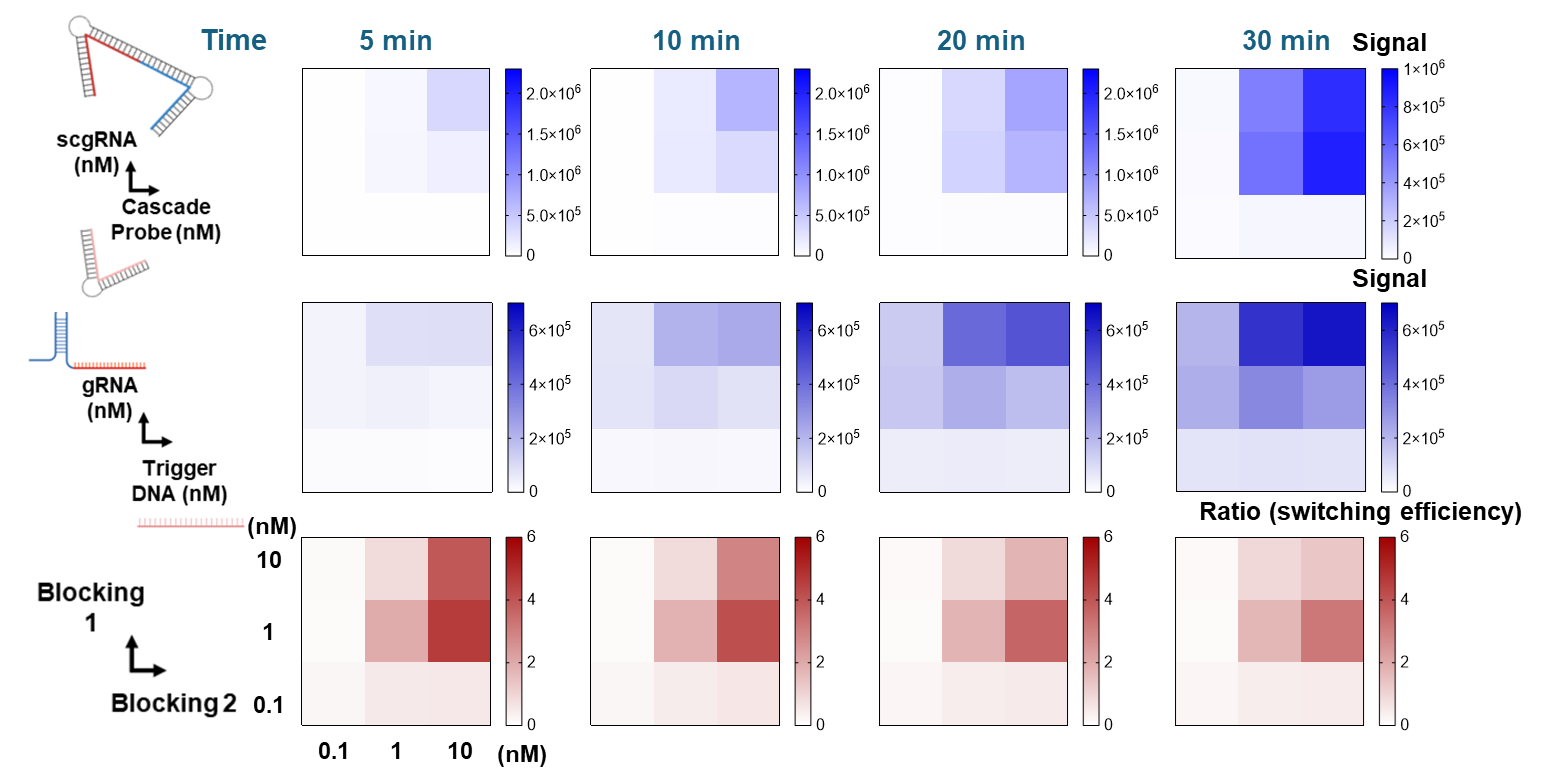

**Figure S9**. Concentration-dependent switching performance of the dual-blocking system. Heat-map matrix screening of scgRNA and Cascade-probe concentrations (0.1, 1, and 10 nM for each component) to identify optimal reaction conditions. The top panel shows fluorescence output when both gRNA and trigger DNA remain blocked, while the middle panel shows signal when both are unblocked. The bottom panel represents the switching ratio (unblocked/blocked) at each concentration pair, serving as a metric for discrimination strength between signal-ON and signal-OFF states. Higher red intensity corresponds to stronger switching performance. Results indicate that 1–10 nM concentrations of both blocking components outperform conditions with only 0.1 nM present. Although 1 nM scgRNA +10 nM Cascade probe yielded the highest switching ratio, a symmetric 10 nM/10 nM configuration was selected for subsequent assays due to comparable performance and ease of implementation.

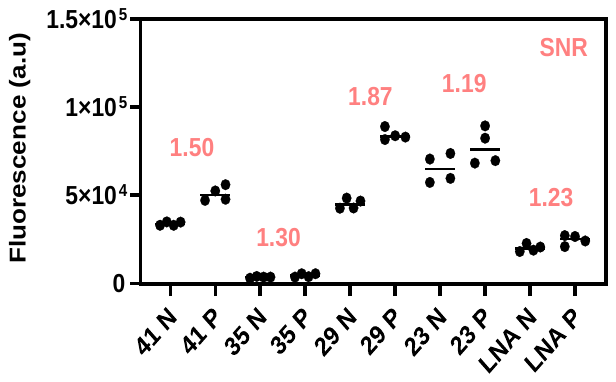

**Figure S10.** End-point fluorescence analysis (t=10 min) of Cascade probe variants in the full Cascade reaction. Cascade probes with varying bubble-DNA lengths and an LNA-modified design (LNA-1) were tested using positive (10⁴ copies/µL MRSA DNA) and negative (no-template) samples. Signal-to-noise ratios (SNRs) were calculated as the ratio of positive (P) to negative (N) signals and are indicated in red for each condition. The 29-nt bubble-DNA variant produced the highest SNR and was selected for subsequent optimization and integration into the dual-blocking system. Data are shown as individual replicates (n = 4).

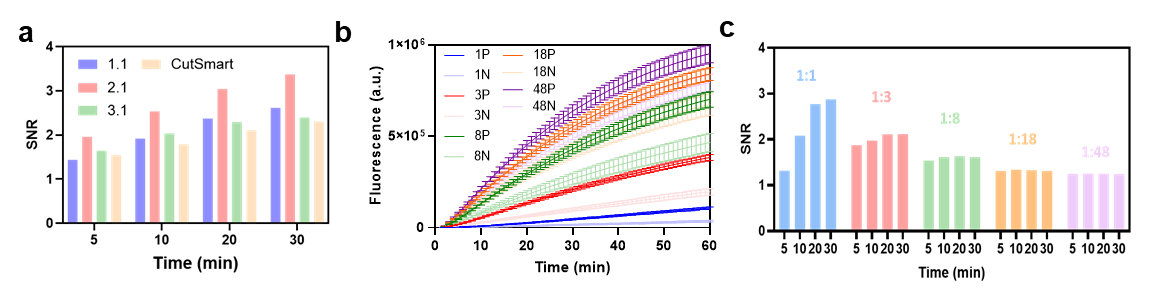

**Figure S11**. Optimization of buffer conditions and T1:T2 ratio for the dual-blocking Cascade assay. (a) End-point fluorescence and kinetic profiles across four reaction buffers (NEB Buffer 1.1, 2.1, 3.1, and CutSmart). NEB 2.1 yielded the highest signal-to-noise ratio (SNR). (b-c) Evaluation of T1:T2 component ratios (1:1, 1:3, 1:8, 1:18, 1:48). As the proportion of T2 increased, fluorescence rose due to a larger pool of autocatalytic components, but the SNR progressively declined and eventually approached unity, making positive and negative reactions indistinguishable. This reflects bypassing of the target dependent T1 activation step, with spontaneous amplification dominating the reaction. These results highlight the need to limit T2 concentration and to optimize the T1:T2 ratio to preserve target specific activation and suppress nonspecific background. The 1:1 ratio produced the strongest SNR; however, a 1:3 ratio was selected for downstream experiments due to improved experimental robustness and ease of distinguishing signal across replicates. Data shown as mean ± s.d. (n = 4).

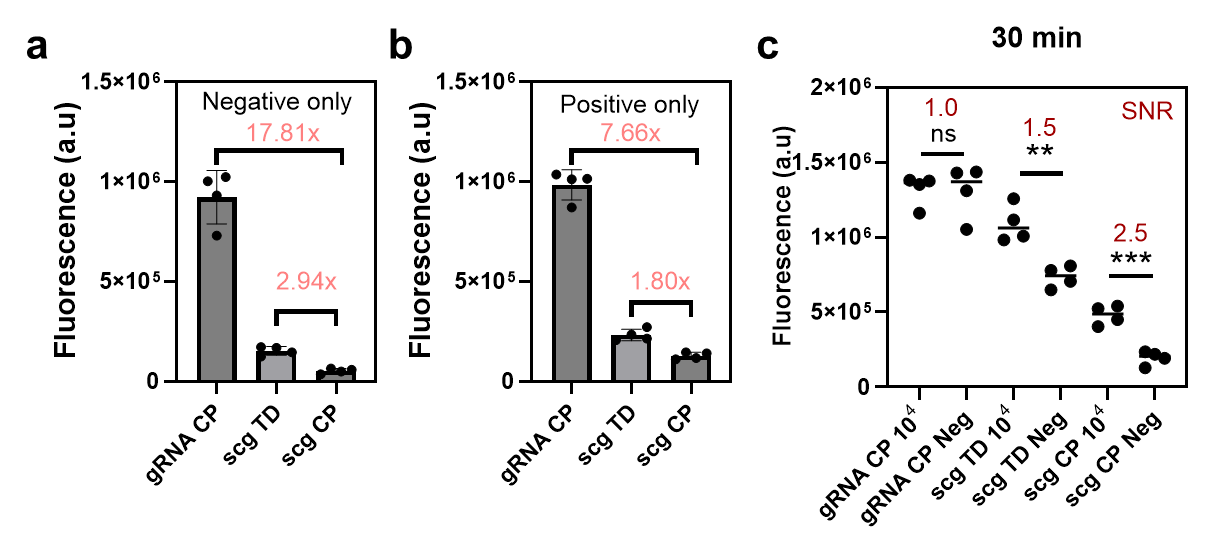

**Figure S12.** Benchmarking dual-blocking versus single-blocking configurations in the full Cascade reaction. (a) Comparison of background leakage under three configurations: dual-blocking (scg CP), scgRNA-only blocking (scg TD), and trigger-DNA-only blocking (gRNA CP). Dual-blocking reduced nonspecific activation by 2.94-fold relative to scgRNA-only blocking and by 17.81-fold relative to trigger-DNA-only blocking. (b) Positive-sample signal comparison under identical conditions showed that dual blocking yielded the lowest signal, with 7.66-fold and 1.80-fold reductions relative to the scgRNA-only and trigger-DNA-only blocking conditions, respectively. (c) End-point signal and SNR analysis at 30 min demonstrate that dual-blocking improves discrimination by disproportionately suppressing leakage. scgRNA-only blocking produced an SNR of 1.5, and trigger-DNA-only blocking yielded an SNR of 1.0, whereas dual-blocking achieved an SNR of 2.5. These data show that although both signal and background decrease under dual-blocking, suppression of spurious activation outweighs loss of true signal, resulting in enhanced SNR. Individual replicates shown (n = 4).

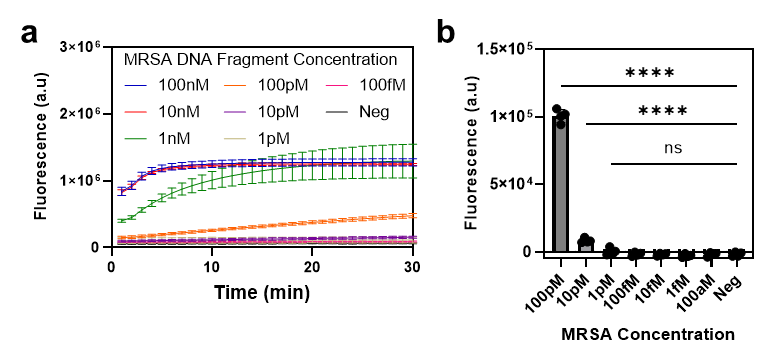

**Figure S13.** Analytical sensitivity of the single-Cas12a RNP T1 reaction. Titration of MRSA DNA fragments in a single-Cas12a (RNP T1-only) configuration demonstrates a detection limit of approximately 10 pM, with diminishing signal at lower concentrations (1 pM–100 aM) approaching negative-control baseline. This result highlights the intrinsic sensitivity limit of non-autocatalytic CRISPR detection and motivates incorporation of the downstream Cascade amplification layer to achieve ultra-sensitive performance. Data shown as mean ± s.d. (n=4).

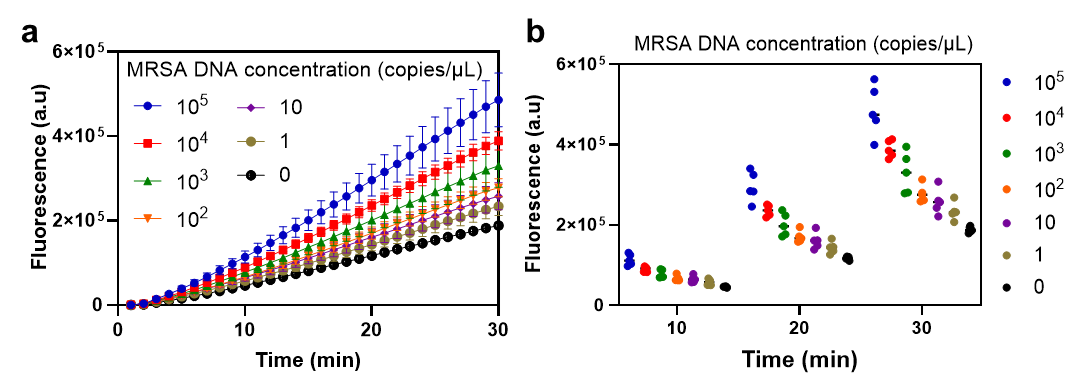

**Figure S14.** Raw fluorescence traces and end-point distribution underlying Fig. 4e. Ten-fold serial dilutions of MRSA genomic DNA (10^5^ to 1 copy/µL) were measured under dual-blocking Cascade conditions using the final protocol of sequential pre incubation followed by pre activation. All individual replicate traces are shown (left), along with the corresponding end-point fluorescence values at 10 min (right), providing the raw data supporting Fig. 4e. As in the main text, signal scaled with input copy number and remained distinguishable down to the single-copy regime. Data are presented as mean ± standard deviation or individual replicate points (n = 5).

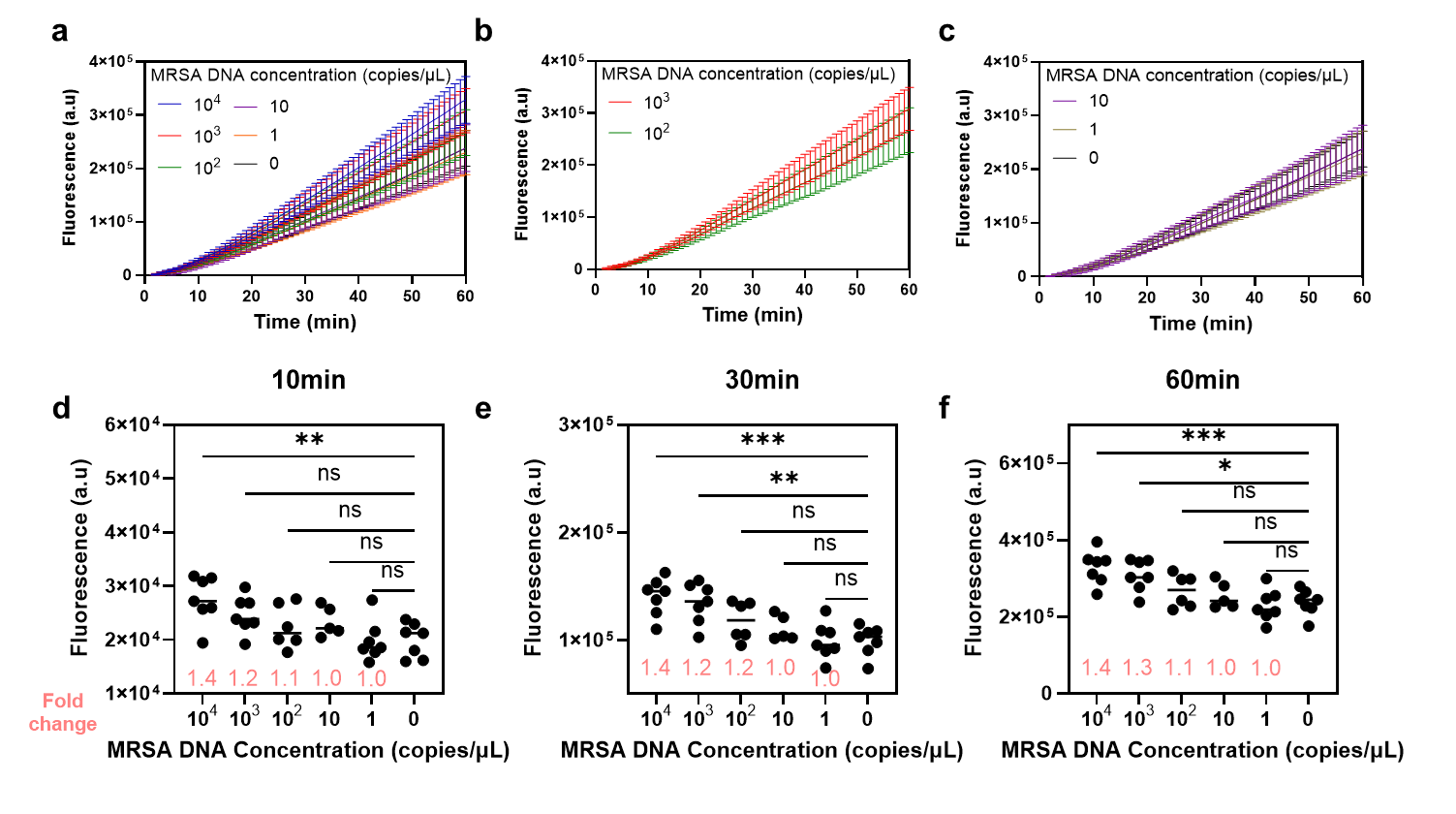

**Figure S15.** Raw fluorescence traces and end-point values for one-pot assembly reactions corresponding to Fig. 4e. All individual replicate data for 10-fold serial dilutions of MRSA genomic DNA (10⁴ to 1 copy/µL) are shown for the one-pot configuration, illustrating loss of discriminatory power below ~10³ copies/µL when RNP T1 and the autocatalytic RNP T2 layer are supplied simultaneously. Data shown as mean ± s.d. or individual replicates (n = 5–7).

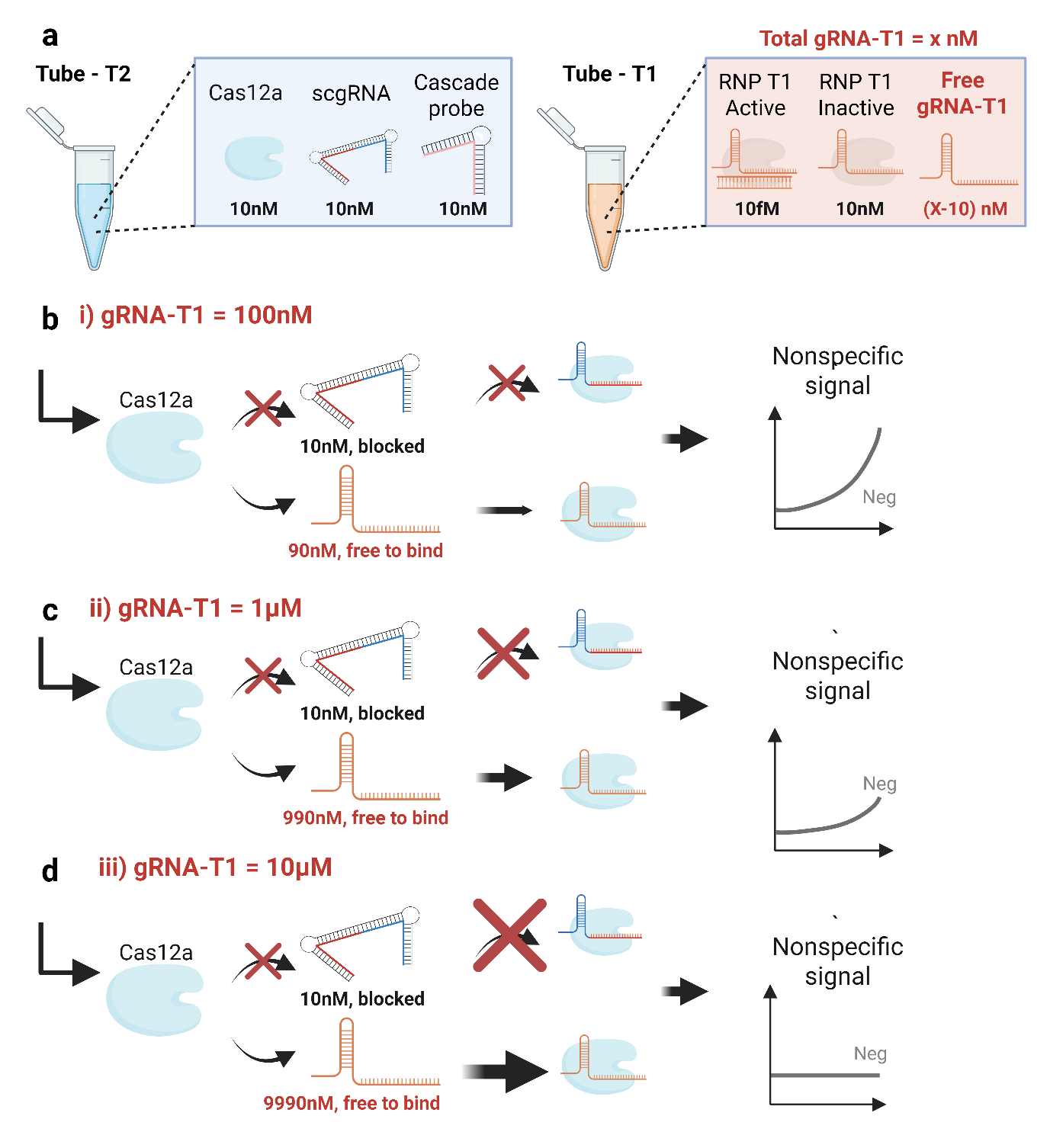

**Figure S16.** Schematic illustrating the competitive gRNA-T1 decoy strategy for leakage suppression.
(a) Tube T1 contained 10 nM Cas12a, *x* nM gRNA-T1, and 10 fM target DNA. After pre-incubation and activation, approximately 10 fM of active RNP T1 and 10 nM of inactive RNP T1 were present, with the 10 fM fraction considered negligible relative to 10 nM. Excess gRNA-T1 therefore remained unbound due to limited Cas12a availability. When the contents of Tube T2 were subsequently introduced, the free Cas12a in Tube T2, uncomplexed with scgRNA, could potentially bind either the blocked scgRNA or the free gRNA-T1. Because of the presumed higher binding affinity and accessibility of the unblocked gRNA-T1, Cas12a preferentially formed RNP complexes with gRNA-T1, effectively sequestering free Cas12a and minimizing nonspecific activation. (b–d) Simulations of three representative cases with increasing total gRNA-T1 concentrations (100 nM, 1 µM, and 10 µM) show that approximately 90 nM, 990 nM, and 9990 nM of gRNA-T1, respectively, remained free after Tube T1 preparation. Upon combining with Tube T2, higher gRNA-T1 concentrations produced stronger suppression of leakage by shifting the binding equilibrium toward the decoy strand and away from premature scgRNA unwinding, thereby reducing nonspecific RNP-T2 activation in a dose-dependent manner.

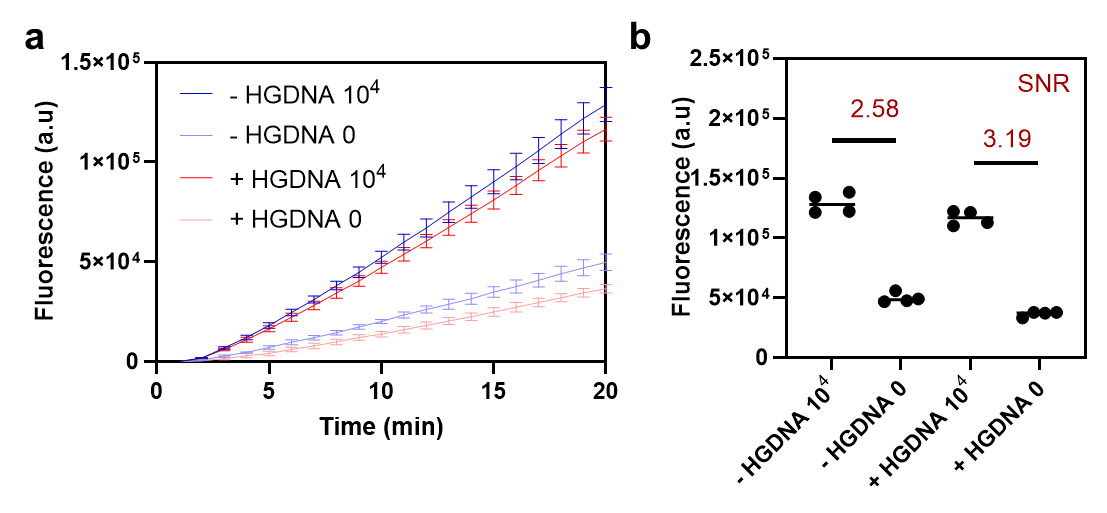

**Figure S17.** Background DNA globally suppresses signal in the dual-blocking Cascade assay. Dual-blocking reactions were supplemented with 3 ng/µL human genomic DNA to assess nonspecific DNA effects on leakage. (a) Fluorescence trajectories show that genomic DNA reduces both positive (10⁴ copies/µL MRSA DNA) and negative signals, indicating global dampening of Cas12a activity rather than selective leakage suppression. (b) SNR improved only modestly (from ~2.58 to ~3.19), consistent with nonspecific sequestration of Cas12a trans-cleavage activity. The attenuation was not pronounced here potentially because the dual-blocking system already minimizes unintended activation and reduces the number of free reactive species. Data displayed as mean ± s.d. or individual replicates (n = 4).

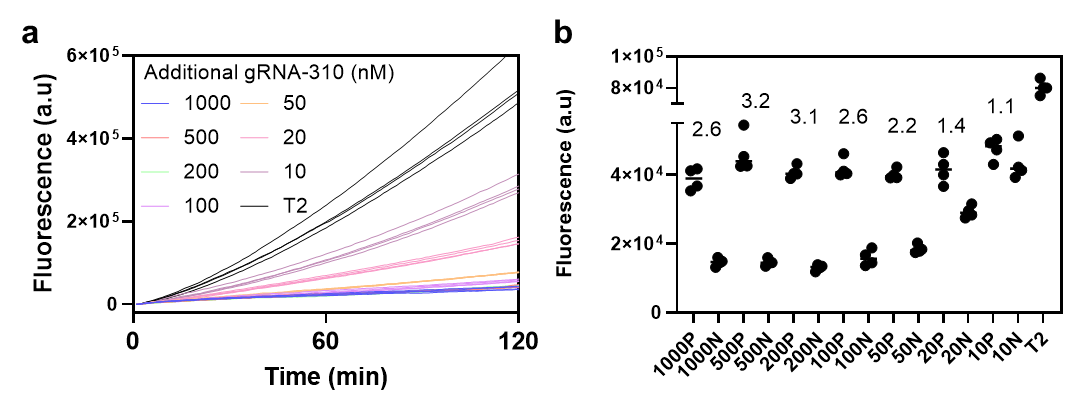

**Figure S18.** Decoy-driven suppression of residual leakage is sequence-independent. Unrelated gRNA decoys (gRNA-310, used in our previous study) exhibited similar background-reduction trends across concentrations ranging from 10 nM to 1 µM. Positive reactions (10⁴ copies/µL MRSA DNA) remained largely unaffected, whereas negative controls showed a dose-dependent decrease in background signal, resulting in improved SNR without compromising true signal. These results confirm that the suppression mechanism based on competitive Cas12a binding by decoy strands is generalizable. Data represent n = 3–4.

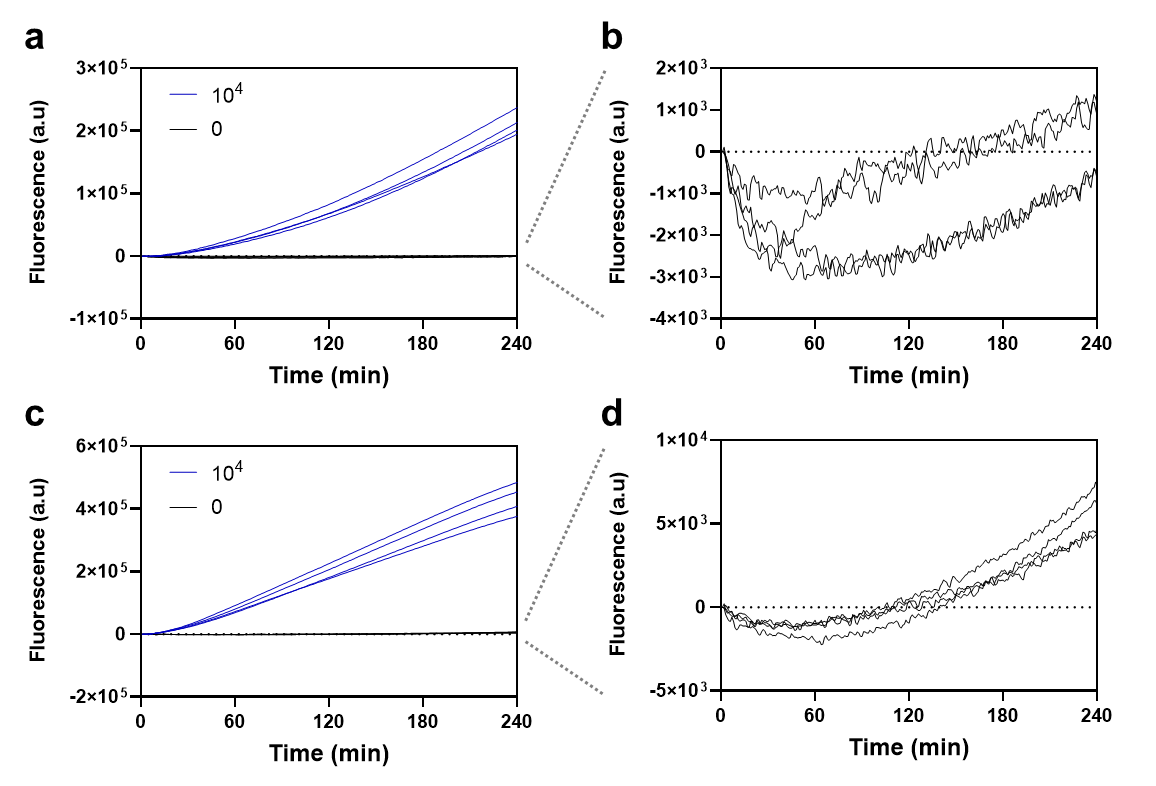

**Figure S19.** (a-b) Extended time-course monitoring of dual-blocking reactions with gRNA-decoy strategy showing that negative-control traces remain flat for several hours, confirming durable leakage suppression, while positive reactions (10⁴ copies/µL MRSA DNA) continue to rise and separate cleanly from negatives, analogous to PCR-style plateau behavior where true negatives remain baseline, demonstrating long-term kinetic stability and sustained discrimination under decoy-assisted conditions (n = 4). (c, d) Technical replicates of (a, b) performed under identical experimental conditions, confirming reproducibility of long-term leakage suppression and signal separation (n = 4).

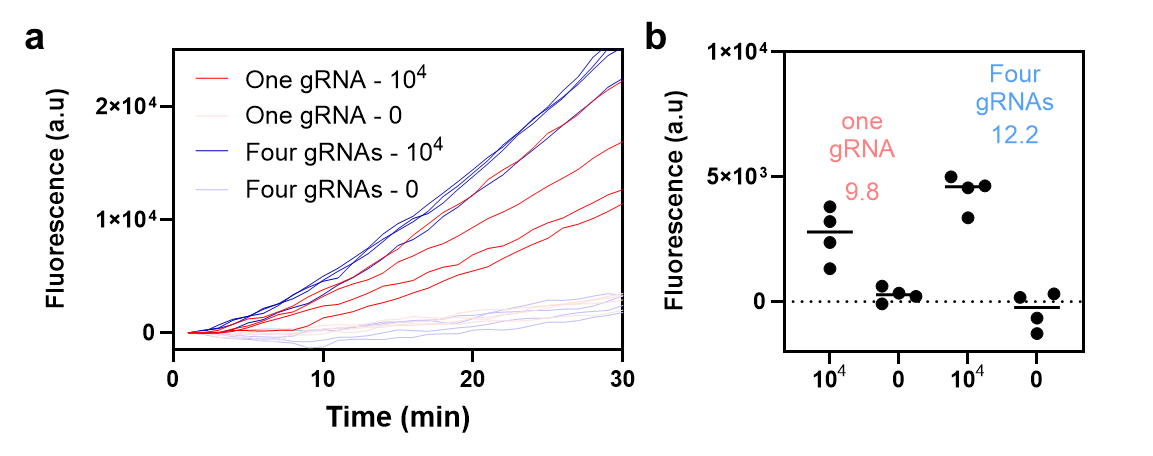

**Figure S20.** Multi-guide enhancement of signal output in the decoy-assisted dual-blocking Cascade system. Fluorescence kinetics for MRSA detection using either a single MRSA-targeting gRNA or a pool of four MRSA-targeting gRNAs (each supplied at 100 nM) show that multi-guide configurations yield stronger signal amplification while maintaining low background. Endpoint signal-to-noise ratios improved from ~9.8 with a single guide to ~12.2 with four guides, demonstrating synergistic activation when multiple target-specific crRNAs are used in parallel. Individual replicate traces are shown (n = 4).

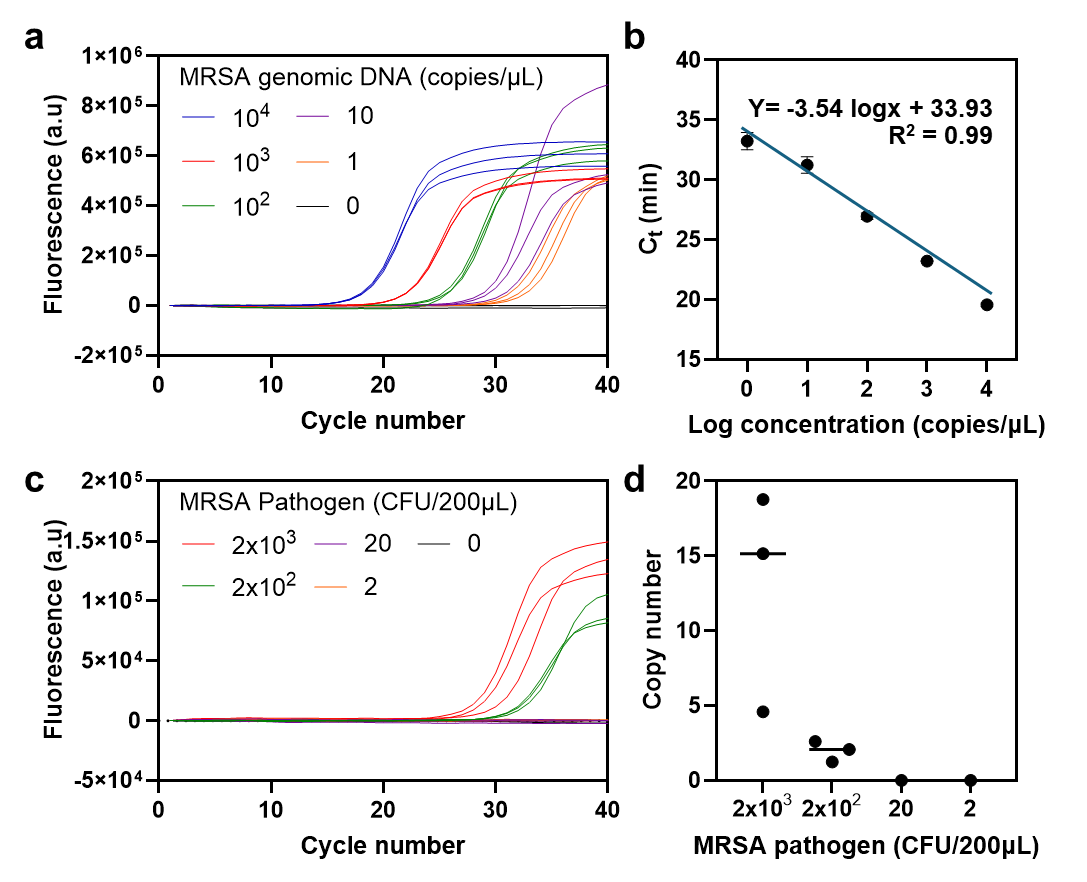

**Figure S21.** Quantification of MRSA DNA extracted from pathogen-spiked whole blood using a commercial kit. (a–b) PCR calibration curve generated with MRSA primers across a range from ten thousand copies per microliter down to one copy per microliter and a zero template control. The assay achieved single-copy sensitivity and showed a clear titration trend with an R-squared value of 0.99. (c–d) PCR results from extracted samples. Only the samples containing 2,000 and 200 CFU/200µL produced detectable signals. Using the calibration curve in panel b, the corresponding copy numbers were back-calculated. The 200 CFU/200µL sample approached the single copy level, and the 20 CFU/200µL sample was not detected. These results highlight the intrinsic limitations of DNA extraction and purification, where binding and release steps inevitably cause loss of target DNA.

**Supplementary Note 1: Varying complement concentration**

For a blocking system with a target ($T$) and complement ($C$) that form an blocked nucleic acid (B$NA$) with an equilibrium:

$$T+C\rightleftharpoons BNA$$

the equilibrium will be described by:

$$K_{eq,1}=\frac{\left[ BNA \right]_{eq}}{\left[ T \right]_{eq}\left[ C \right]_{eq}}$$

We can solve exactly for $\left[ T \right]_{eq}$ for some initial concentrations (e.g. $\left[ T \right]_{0}=10 nM, \left[ C \right]_{0}=15 nM$) and some typical $K_{eq}$:

$$K_{eq}=\frac{\left[ BNA \right]_{eq}}{\left[ T \right]_{eq}\left[ C \right]_{eq}}=\frac{\left[ T \right]_{0}-\left[ T \right]_{eq}}{\left[ T \right]_{eq}\left( \left[ C \right]_{0}-\left[ BNA \right]_{eq} \right)}=\frac{\left[ T \right]_{0}-\left[ T \right]_{eq}}{\left[ T \right]_{eq}\left( \left[ C \right]_{0}-\left[ T \right]_{0}+\left[ T \right]_{eq} \right)}$$

This results in a quadratic equation:

$${K_{eq}\left[ T \right]}_{eq}^{2}+\left[ 1+K_{eq}\left( \left[ C \right]_{0}-\left[ T \right]_{0} \right) \right]\left[ T \right]_{eq}-\left[ T \right]_{0}=0$$

With the positive solution:

$$\left[ T \right]_{eq}=\frac{-\left[ 1+K_{eq}\left( \left[ C \right]_{0}-\left[ T \right]_{0} \right) \right]+\sqrt{\left[ 1+K_{eq}\left( \left[ C \right]_{0}-\left[ T \right]_{0} \right) \right]^{2}+4K_{eq}\left[ T \right]_{0}}}{2K_{eq,1}}$$

$$=\frac{K_{eq}\left( \left[ T \right]_{0}-\left[ C \right]_{0} \right)-1+\sqrt{K_{eq}^{2}\left( \left[ C \right]_{0}-\left[ T \right]_{0} \right)^{2}+2K_{eq}\left( \left[ C \right]_{0}+\left[ T \right]_{0} \right)+1}}{2K_{eq}}$$

We can estimate $K_{eq}$ from the change in free energy for hybridization. For example, for $\Delta G\approx-15 kcal/mol$ at 33˚C:

$$K_{eq}=e^{{-\Delta G}/{RT}}\approx5\times{10}^{10}$$

A table of solutions reveals the effect of changing the concentration of the complement on the equilibrium concentration of target:

| $\left[ \boldsymbol{T} \right]_{\boldsymbol{0}}\boldsymbol{(nM)}$ | $\left[ \boldsymbol{C} \right]_{\boldsymbol{0}}\boldsymbol{(nM)}$ | $\left[ \boldsymbol{C} \right]_{\boldsymbol{0}}\boldsymbol{:}\left[ \boldsymbol{T} \right]_{\boldsymbol{0}}$ | $\left[ \boldsymbol{T} \right]_{\boldsymbol{eq}}$ |
| --- | --- | --- | --- |
| 10 | 10 | 1:1 | 430 pM |
| 10 | 15 | 1:1.5 | 38.7 pM |
| 10 | 20 | 1:2 | 19.5 pM |
| 10 | 100 | 1:10 | 2.18 pM |
| 10 | 1000 | 1:100 | 198 nM |

This effect also depends on the value of $\Delta G$:m
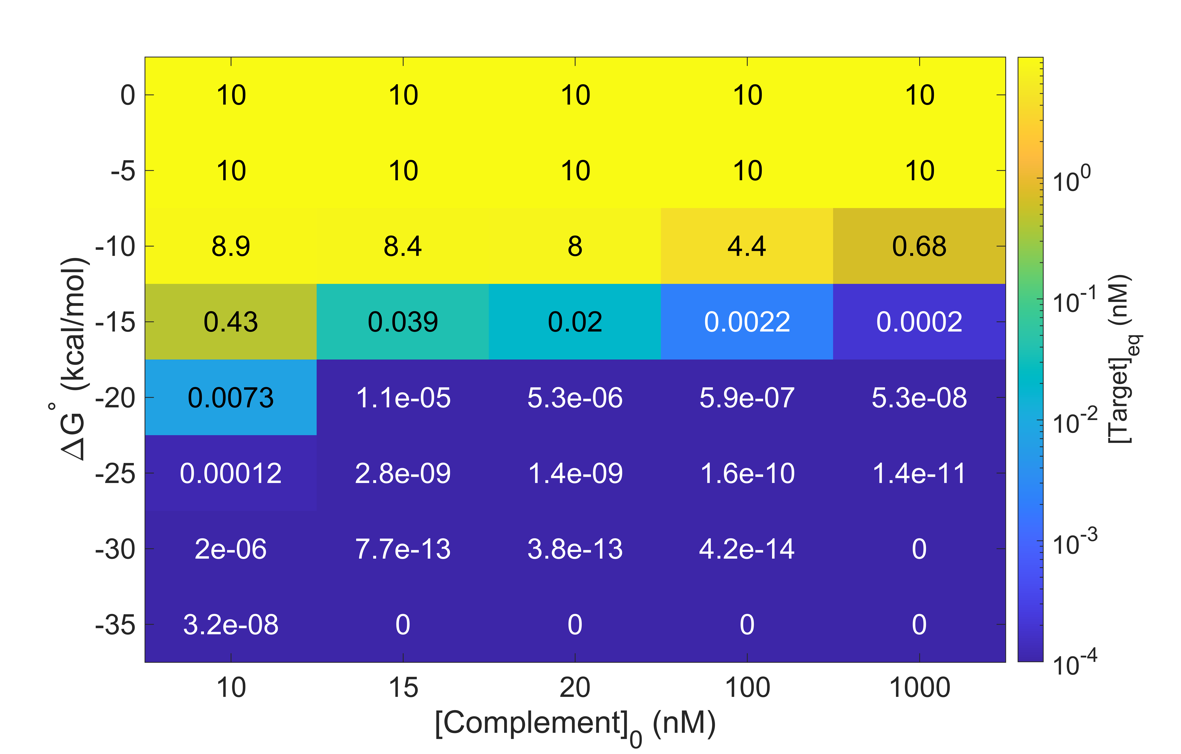

For high $\Delta G$ values, nanomolar concentrations are not sufficient to drive the reaction forward significantly. For low $\Delta G$ values, however, the concentration of target is negligible. Therefore, the equation:

$$\left[ T \right]_{eq}=\frac{\left[ BNA \right]_{eq}}{K_{eq}\left[ C \right]_{eq}}$$

can be approximated by:

$$\left[ T \right]_{eq}=\frac{\left[ T \right]_{0}-\left[ T \right]_{eq}}{K_{eq}\left( \left[ C \right]_{0}-\left[ T \right]_{0}+\left[ T \right]_{eq} \right)}\approx\frac{\left[ T \right]_{0}}{K_{eq}\left( \left[ C \right]_{0}-\left[ T \right]_{0} \right)}$$

Meaning $\left[ T \right]_{eq}$ is inversely proportional to the excess concentration of the complement:

$$\left[ T \right]_{eq}\approx\frac{\left[ T \right]_{0}}{K_{eq}\left[ C \right]_{0,excess}}$$

**Supplementary Note 2: Double blocking rationale**

If we expand to two blocking systems, we have two equilibria:

$$T_{1}+C_{1}\rightleftharpoons BNA_{1}$$

$$T_{2}+C_{2}\rightleftharpoons BNA_{2}$$

We can consider a system where both targets are required to activate the enzyme, such as a gRNA and ssDNA target for Cas12. In this case, the RNP complex formation is a simple equilibrium:

$$E+T_{1}\rightleftharpoons ET_{1} \text{(RNP complex formation)}$$

This reaction is described by:

$$K_{eq}=\frac{\left[ ET_{1} \right]_{eq}}{\left[ E \right]_{eq}\left[ T_{1} \right]_{eq}}$$

The later step activating the enzyme can be modeled as:

$$ET_{1} +T_{2}\rightleftharpoons\left( ET_{1} \right)T_{2}\to E_{act} \text{(cis-cleavage)}$$

This includes an irreversible catalytic step because cis-cleavage involves separate binding and cleavage events. Because the enzyme is consumed, however, this cannot be treated as a Michaelis-Menten reaction and the initial velocity is not easily approximated. To simplify, we can reduce the reaction to:

$$ET_{1} +T_{2}\to E_{act}$$

The rate of active enzyme production will then dictate the speed at which the autocatalytic cycle is initiated, causing nonspecific signal. Initially, this reaction proceeds according to the rate:

$$v=k_{cis}\left[ ET_{1} \right]_{eq}\left[ T_{2} \right]_{eq}$$

The species conservation for the enzyme is:

$$\left[ E \right]_{0}=\left[ E \right]+\left[ ET_{1} \right]+\left[ E_{act} \right]$$

Early in the reaction, when $\left[ E_{act} \right]$ is small, we can simplify to:

$$\left[ E \right]_{0}=\left[ E \right]+\left[ ET_{1} \right]$$

To replace $\left[ ET_{1} \right]_{eq}$:

$$\left[ ET_{1} \right]_{eq}=\left[ E \right]_{0}-\left[ E_{1} \right]_{eq}=\left[ E \right]_{0}-\frac{\left[ ET_{1} \right]_{eq}}{K_{eq}\left[ T_{1} \right]_{eq}}$$

$$\left[ ET_{1} \right]_{eq}={\left( \frac{K_{eq}\left[ T_{1} \right]_{eq}}{1+K_{eq}\left[ T_{1} \right]_{eq}} \right)\left[ E \right]}_{0}$$

Substituting in the equation for $v$:

$$v=k_{cis}\left[ ET_{1} \right]_{eq}\left[ T_{2} \right]_{eq}={\left( \frac{k_{cis}K_{eq}\left[ T_{1} \right]_{eq}\left[ T_{2} \right]_{eq}}{1+K_{eq}\left[ T_{1} \right]_{eq}} \right)\left[ E \right]}_{0}$$

The parameter $\alpha_{1}$ is defined as the fraction of target molecules in the annealed state at equilibrium:

$$\alpha_{x}=\frac{\left[ BNA_{x} \right]_{eq}}{\left[ T_{x} \right]_{0}}=\frac{\left[ BNA_{x} \right]_{eq}}{\left[ T_{x} \right]_{eq}+\left[ BNA_{x} \right]_{eq}}$$

where $\left[ T_{x} \right]_{0}$ is the amount of target initially added to the system.

Consequently:

$$\left[ BNA_{x} \right]_{eq}=\alpha_{x}\left[ T_{x} \right]_{0}$$

$$\left[ T_{x} \right]_{eq}=\left( 1-\alpha_{x} \right)\left[ T_{x} \right]_{0}$$

This parameter is useful to describe the tendency of the BNA to remain bound under certain reaction conditions (as an alternative to $K_{eq}$). Because $\alpha$ relates directly to the amount of unbound target at equilibrium, it can provide insight into the amount of nonspecific signal caused by the thermodynamic equilibrium of a certain SNA.

In terms of $\alpha$, the velocity can be stated as:

$$v=\frac{k_{cis}K_{eq}\left( 1-\alpha_{1} \right)\left( 1-\alpha_{2} \right)\left[ T_{1} \right]_{0}\left[ T_{2} \right]_{0}}{1+K_{eq}\left( 1-\alpha_{1} \right)\left[ T_{1} \right]_{0}}\left[ E \right]_{0}$$

This numerator indicates that the suppression of noise caused by the respiratory effect is a product of both $\alpha$ values. If only one gate was functioning ($\alpha_{1}=0.999, \alpha_{2}=0)$, the $\left( 1-\alpha_{1} \right)\left( 1-\alpha_{2} \right)$ term is $0.001$. If both $\alpha$ are $0.999$, on the other hand, the value of the term is $0.000001$. This is the key numerical justification of the double blocking approach.

The behavior of this equation can be further classified depending on the denominator. Sinan et al. reported a $K_{d}$ for Cas12a RNP formation of $0.6 pM$, or $K_{eq}\approx1.67\times{10}^{12} M^{-1}.$^1^ For a typical target concentration like $10 nM$, the term $K_{eq}\left( 1-\alpha_{1} \right)\left[ T_{1} \right]_{0}$ is approximately ${10}^{4}\left( 1-\alpha_{1} \right)$. For small $\alpha_{1}$ (poorly functioning first gate), ${10}^{4}\left( 1-\alpha_{1} \right)\gg1$. This term dominates the denominator, resulting in:

$$v\approx k_{cis}\left( 1-\alpha_{2} \right)\left[ T_{2} \right]_{0}\left[ E \right]_{0}$$

so, the rate is determined solely by the function of the second gate. A poorly functioning second gate ($\alpha_{2}\approx0)$, on the other hand, will result in:

$$v\approx\frac{k_{cis}K_{eq}\left( 1-\alpha_{1} \right)\left[ T_{1} \right]_{0}\left[ T_{2} \right]_{0}}{1+K_{eq}\left( 1-\alpha_{1} \right)\left[ T_{1} \right]_{0}}\left[ E \right]_{0}$$

Comparing these as a ratio of $v$ values:

$$\frac{v_{poor \alpha_{2}}}{v_{poor \alpha_{1}}}=\frac{K_{eq}\left( 1-\alpha_{1} \right)\left[ T_{1} \right]_{0}}{\left( 1+K_{eq}\left( 1-\alpha_{1} \right)\left[ T_{1} \right]_{0} \right)\left( 1-\alpha_{2} \right)}\approx\frac{1}{1-\alpha_{2}}$$

This indicates that similarly poor performance of the second gate contributes more strongly to nonspecific signal when compared to poor performance of the first gate.

When the first gate is functioning very well ($\alpha_{1}\approx1,\left( 1-\alpha_{1} \right)\ll{10}^{-4}$), the ${10}^{4}\left( 1-\alpha_{1} \right)$ term has little effect on the velocity and the equation simplifies to:

$$v=k_{cis}K_{eq}\left( 1-\alpha_{1} \right)\left( 1-\alpha_{2} \right)\left[ T_{1} \right]_{0}\left[ T_{2} \right]_{0}\left[ E \right]_{0}$$

If we assume that both gates function similarly ($\alpha_{1}=\alpha_{2}$ and $\left[ T_{1} \right]_{0}=\left[ T_{2} \right]_{0}$), we can simplify to:

$$v=\frac{k_{cis}K_{eq}\left( 1-\alpha\right)^{2}\left[ T \right]_{0}^{2}}{1+K_{eq}\left( 1-\alpha\right)\left[ T \right]_{0}}\left[ E \right]_{0}$$

For well-functioning gates, this further simplifies to:

$$v=k_{cis}K_{eq}\left( 1-\alpha\right)^{2}\left[ T \right]_{0}^{2}\left[ E \right]_{0}$$
